# Dissecting the eQTL micro-architecture in *Caenorhabditis elegans*

**DOI:** 10.1101/651885

**Authors:** Mark G. Sterken, Roel P. J. Bevers, Rita. J. M. Volkers, Joost A. G. Riksen, Jan E. Kammenga, L. Basten Snoek

## Abstract

The study of expression quantitative trait loci (eQTL) using natural variation in inbred populations has yielded detailed information about the transcriptional regulation of complex traits. Studies on eQTL using recombinant inbred lines (RILs) led to insights on local and distant regulatory loci of transcript abundance. However, determining the underlying causal polymorphic genes or variants is difficult, but ultimately essential for the understanding of regulatory networks of complex traits. This requires insight into whether associated loci are single eQTL or a combination of closely linked eQTL, and how this QTL micro-architecture depends on the environment. We addressed these questions by mapping eQTL in N2 x CB4856 *C. elegans* RIL populations across three different environments (control, heat-stress, and recovery). To test for independent replication of the RIL eQTL, we used introgression lines (ILs). Both populations indicate that the overall heritability, number, and position of eQTL differed among environments. Across environments we were able to replicate 70% of the local- and 40% of the distant-eQTL using the ILs. Simulation models revealed that additive effects explain up to 60-93% of RIL/IL heritability across environments. Closely linked eQTL explained up to 40% of RIL/IL heritability in the control environment whereas only 7% in the heat-stress and recovery environments. In conclusion, we show that reproducibility of eQTL was higher for local vs. distant eQTL and that the environment affects the eQTL micro-architecture.

## Introduction

The genetic architecture of quantitative traits in genetically segregated populations such as recombinant inbred lines (RILs) differs within and across species and strongly depends on the environment and type of trait (Fournier and Schacherer 2017). In some cases, quantitative traits are (nearly) monogenic and display Mendelian characteristics. For instance the Kallman syndrome in humans is caused by a knockout of FGFR1 (which encodes fibroblast growth factor receptor 1) (Muenke *et al.* 1994), industrial melanism in British peppered moths is caused by a single gene mutation (Van’t Hof *et al.* 2011), and natural variation in the *npr-1* gene in the nematode *Caenorhabditis elegans* is a major determinant of multiple life-history traits (Andersen *et al.* 2014; Sterken *et al.* 2015). These monogenic traits are relatively easy to detect but rather exceptions than the rule. Most quantitative traits display a complex polygenic genetic architecture, regulated by multiple QTLs, where each locus explains a proportion of the trait variation (Albert and Kruglyak 2015).

Studying quantitative genetics of the transcriptome has proven useful in understanding the architecture of complex traits. Expression quantitative trait loci (eQTL) can be measured reliably in a high-throughput fashion in many organisms like: yeast, *Arabidopsis thaliana*, tomato, maize, *C. elegans*, mice, and humans (Schadt *et al.* 2003; Brem and Kruglyak 2005; Li *et al.* 2006; Keurentjes *et al.* 2007; Ranjan *et al.* 2016). The advantage of the eQTL-approach lies in the assessment of thousands of traits simultaneously for which the genetic architectures can be compared (Jansen and Nap 2001; Gilad *et al.* 2008). First, the variance explained per underlying QTL will cover the whole range from essentially monogenic (Mendelian) to nearly undetectable. Second, correlating gene expression to loci can reveal the biological processes and phenotypes that lay downstream of (e)QTL (*e.g.* (Jimenez-Gomez *et al.* 2010; Terpstra *et al.* 2010; Jimenez-Gomez *et al.* 2011; Andersen *et al.* 2014; Schmid *et al.* 2015; Sterken *et al.* 2017; Albert *et al.* 2018)). In general, two patterns stand out in eQTL studies: i) the occurrence of regulatory hotspots (*trans*-bands) where a single locus affects the expression of many genes across the genome and ii) an abundance of locally regulated eQTL (*cis*-eQTL) where the locus is located on or near the gene of which it affects the expression (Nica and Dermitzakis 2013; Albert *et al.* 2018). *Cis*-eQTL can occur due to polymorphisms affecting the transcription of the gene under study, *e.g.* deletions or non-functional promotor regions. The occurrence of *trans*-bands was found to be specific for environmental conditions linked to genetic effects. Together, the *cis*- eQTL and *trans*-bands typically cover ∼75% of the mapped eQTL in any study (e.g. (Snoek *et al.* 2017; Albert *et al.* 2018)).

The detection of (e)QTL depends on many different factors like developmental stage, background mutations, and the ambient environment, to name a few (Li *et al.* 2006; Vinuela *et al.* 2010; Duveau and Felix 2012; Snoek *et al.* 2012; Andersen *et al.* 2014; Francesconi and Lehner 2014). Importantly, QTL detection is mainly determined by the number of genetically segregating RILs and the density and structure of the genetic marker map. Further characterization of QTLs often involves the identification of the underlying causal polymorphic genes in order to understand molecular mechanisms affecting complex traits and/or to facilitate breeding for selected traits. An important hallmark of QTL mapping has been the fine mapping and identification of polymorphic regulators or quantitative trait nucleotides (QTN) (Radwan and Babik 2012; Rockman 2012).

Fine mapping of causal variants requires determining if associated loci are single QTL or combinations of closely linked QTLs, here coined the QTL micro-architecture, and how this micro-architecture depends on the environment. Apparent single QTLs may be truly single QTL or a combination of closely linked QTLs not segregating in the studied population. These are often hard to characterize because of a lack of resolution of the genetic map or shortage of a sufficient number of recombination events. Here, we set out to investigate the QTL micro-architecture of eQTLs in the nematode *C. elegans* in a segregated population of RILs derived from a cross between wild-types Bristol N2 and CB4856 (Li *et al.* 2006; Thompson *et al.* 2015). We use a set of previously detected eQTL for their replication in a separate, independent, population of introgression lines (ILs) in this study (Doroszuk *et al.* 2009; Thompson *et al.* 2015; Snoek *et al.* 2017). Compared to the RILs, which are genetic mosaics of loci derived from both parents, ILs contain a single genomic segment of one parent in a genetic background of another parent. Typically, introgression lines are used to verify the existence of QTL, or to narrow down the location of a QTL in order to find the underlying causal polymorphism (*e.g.* see (Gao *et al.* 2018)). The RIL and IL populations are exposed to three environments: heat-shock, recovery from heat-shock, and a control environment at a standard rearing temperature. We investigate three aspects of eQTL mapped in RILs: i) we address how well eQTL are replicated in ILs, ii) we test if the predicted numbers of eQTL agree with the observed number of differentially expressed genes in ILs, and iii) we simulate QTL architectures explaining the relative heritability in RILs and ILs and match our data to these models. Finally, we show that the eQTL microarchitecture under ambient conditions consists of closely linked eQTL.

## Material and methods

### Strains used

The wild-type strains N2 and CB4856 were used and 57 introgression lines (ILs) with segments of CB4856 in an N2 genetic background (Doroszuk *et al.* 2009). Most of the ILs have been sequenced, and the genetic map was based on the sequenced genotypes (Thompson *et al.* 2015), a file with the strains and the map has been included in **Table S1**. The data used for the recombinant inbred lines (RILs) is accessible in (Snoek *et al.* 2017).

### Nematode culturing

The strains were cultured as described previously (Snoek *et al.* 2017). In short, strains were kept in maintenance at 12 °C on 6-cm Nematode Growth Medium (NGM) Petri dishes seeded with the *Escherichia coli* strain OP50 as food source (Brenner 1974). Before starting the experiment, single hermaphrodites of each strain in the L2 stage were sub-cultured in 12-wells plates and grown at 20 °C. The offspring was screened for the occurrence of males by microscopy and only populations consisting solely of hermaphrodites were transferred to 9-cm NGM Petri dishes containing *E. coli* OP50 and grown until the bacterial food was depleted.

### Treatments for the transcriptomic experiment

The experiment described in our previous paper was repeated on the IL population and the parental strains (Snoek *et al.* 2017). We started the experiment by transferring a starved population to a new 9-cm NGM Petri dish seeded with *E. coli* OP50. After 60 h at 20 °C, the populations consisting of egg-laying adults were bleached to obtain the eggs, which were transferred to a new 9-cm NGM Petri dish (Brenner 1974). Around 46 h the developmental stage of the population was assessed by microscopy, verifying that the population consisted mainly of L4 larvae. Populations not consisting of L4 larvae were not used. The strains were exposed to one of three environments: (i) a control environment; grown for 48 h at 20 °C, (ii) a heat-stress environment; grown for 46 h at 20 °C and subsequently exposed to 35 °C for 2 h, and (iii) a recovery environment; grown for 46 h at 20 °C, exposed to 35 °C for 2 h, and thereafter returned to 20 °C for 2 h. Directly after the treatment the animals were washed off the Petri dish using M9 buffer and collected in a 1.5 mL Eppendorf tube, which was centrifuged and the pellet was flash frozen in liquid nitrogen. After quality control of the RNA isolation, we obtained 56 ILs for the control and heat-stress environment, and 55 ILs for the recovery environment. For the parental strain N2: four control samples, four heat stress samples and three recovery samples passed quality control. For the parental strain CB4856: three control samples, three heat stress samples and five recovery samples passed quality control.

### RNA isolation

The procedure was followed as described before (Snoek *et al.* 2017). The RNA of the samples was isolated using the RNEasy Micro Kit from Qiagen (Hilden, Germany) following the ‘Purification of Total RNA from Animal and Human Tissues’. The lysis step was modified, pellets were lysed in 150 µl RLT buffer, 295 µl RNAse-free water, 800 µg/ml proteinase K and 1% ß-mercaptoethanol, which was incubated at 55 °C and 1000 rpm in a Thermomixer (Eppendorf, Hamburg, Germany) for 30 minutes, where after the manufacturer’s protocol was followed.

### cDNA synthesis, labelling and hybridization

Before starting cDNA synthesis, the quality and quantity of the RNA was measured by NanoDrop-1000 spectrophotometer (Thermo Scientific, Wilmington DE, USA). The integrity of the RNA was assessed by loading 3 μL per sample on a 1% agarose gel. Samples with good quality RNA were processed following the ‘Two-Color Microarray-Based Gene Expression Analysis; Low Input Quick Amp Labeling’ -protocol, version 6.0 from Agilent (Agilent Technologies, Santa Clara, CA, USA). The microarrays used were Agilent *C. elegans* (V2) Gene Expression Microarray 4X44K slides.

### Array scanning, data extraction, and normalization

After washing, the microarrays were scanned using an Agilent High Resolution C Scanner following the recommended settings. For data extraction, Agilent Feature Extraction Software (version 10.7.1.1) was used, following manufacturer’s guidelines. The extracted data was normalized using the limma package in “R” (version 3.4.2, x64) (Ritchie *et al.* 2015). As recommended, the data was not background corrected before normalization (Zahurak *et al.* 2007), the within-array normalization used the ‘Loess’ method and between-array normalization used the Quantile method (Smyth and Speed 2003).

### Data analysis

Data was analyzed using “R” (version 3.4.2, x64) with custom written scripts (Team 2017), accessible via https://git.wur.nl/published_papers/sterken_2019_closely_linked_qtl. For analysis, the dplyr and tidyr packages were used for data organization (Wickham 2018a; Wickham 2018b), and plots were generated using ggplot2 (Wickham 2009), except for plots displaying simulation results. The transcriptome data analyzed was deposited at ArrayExpress under E-MTAB-5779 (RIL data) (Snoek *et al.* 2017), and E-MTAB-7424 (IL and parental data, described in this paper).

### eQTL data

The eQTL data from the RILs were obtained from a previous publication (Snoek *et al.* 2017) and can be explored at www.bioinformatics.nl/wormqtl2 (Snoek *et al.* 2019). In short, eQTL were mapped using a single marker model, which was conducted separately for each environment. The threshold for each of the three environments was set at –log_10_(p) > 3.9 (false discovery rate, FDR = 0.05) (see (Snoek *et al.* 2017) for details).

### Data transformation

The expression data was transformed to a z-score based on the standard deviation and mean of the N2-replicates per environment, using

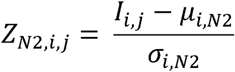

where *Z_N2_* is the Z-score of spot *i* (1, 2, …, 45220) of strain *j* (one of the RILs or ILs) and *I* is the normalized intensity. The *µ* is the mean of the intensity over the N2 strains for spot *i* and the σ is the standard deviation over the N2 strains for spot *i*. This transformation shows the effect of the CB4856 introgressions in the ILs.

Additionally, the expression data was transformed to a z-score based on the standard deviation and mean of the CB4856-replicates per environment, using

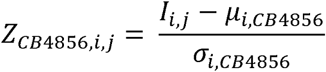

where *Z_CB4856_* is the Z-score of spot *i* (1, 2, …, 45220) of strain *j* (one of the RILs or ILs) and *I* is the normalized intensity. The *µ* is the mean of the intensity over the CB4856 strains for spot *i* and the σ is the standard deviation over the CB4856 strains for spot *i*. This transformation shows the effect of the genetic background in the ILs.

In order to compare IL gene expression with eQTL effect sizes, another data transformation was used, as there the number of standard deviations difference with the mean is less informative than the effect in log_2_-ratio with the mean. Hence, we transformed the data by

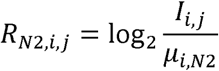

where *R_N2_* is the log_2_-ratio with the mean of spot *i* of strain *j*, *I* is the normalized intensity, and *µ* is the mean intensity over the N2 samples.

### Confirmation of IL genotype

Before mapping, we assured that samples were labelled correctly by applying a *cis*-eQTL based analysis of gene expression compared to the genetic map of the IL population, for details and explanation, see (Zych *et al.* 2017). Samples of three strains were removed (WN263, WN264, and WN284), as the strains did not match the genotypes in the IL population (Doroszuk *et al.* 2009; Thompson *et al.* 2015).

### Confirmation of eQTL in introgression lines

The differential expression per IL was calculated based on the Z-score calculated using the N2 parental strains (Z_N2_). As each IL was measured once, we estimated the significance assuming a normal distribution based on the genetic-background parent. These significances were corrected for multiple testing by the Benjamini-Hochberg method, as applied by the *p.adjust* function in R (Benjamini and Hochberg 1995). As a control and benchmark, the same procedure was applied to the individual RILs. The number of differentially expressed genes with an expected eQTL per CB4856 locus were counted for both the ILs and the RILs. These were expressed as a percentage of the expected eQTL in that locus (separately for *cis*- and *trans*-eQTL).

Another method of confirming eQTL is by correlating the differential expression per inbred line compared to the genetic background-parent (R_N2_) with the QTL effect. By calculating the Pearson correlation of R_N2_ with the eQTL effect per IL or RIL separately for the *cis*- and *trans*-eQTL, it can be assessed how well the overall QTL patterns were recapitulated per strain. These correlations were calculated using the *cor* function and the significances of correlation were calculated using the *cor.test* function.

### Confirmation of trans-bands using ILs

In the eQTL study on the RILs (Snoek *et al.* 2017), 19 *trans*-bands were identified (**Table S3**). For each *trans*-band 2 to 6 ILs covering or flanking the locus were tested by correlating the *trans*-eQTL effect with the R_N2_ in the ILs. Significance was determined based on the correlation values in the N2 samples. Only if the correlation was stronger than the highest-correlating N2 sample, it was called as significant. Only significant positive correlations (effect direction matches the eQTL model) were scored as confirming the *trans*-band.

To determine the overall false-positive rate of this analysis, we determined the correlation of the ILs that were not in or near the *trans*-band region. These correlations were compared to the correlation value at which the *trans*-band was called confirmed. This resulted in an overall false positive rate of 0.11. It should be noted that this approach assumes that *trans*-bands stem from a single location and that these locations were accurately pinpointed in the RIL mapping. Hence, the derived threshold can be considered strict.

### Expectation of differential expressed genes based on eQTL

First, the number of differentially expressed genes in each RIL and IL was calculated using the contrast with either of the ancestral strains, Z_N2_ and Z_CB4856_, at an estimated significance of p < 1*10^-5^. Second, the expected number of differentially expressed genes was estimated for each RIL and IL based on the eQTL architecture in the RIL population (eQTL threshold, - log_10_(p) > 3.9; FDR = 0.05) (Snoek *et al.* 2017). These values were normalized to the RIL population (the RIL population average was set to 1), to compare the relative number of differentially expressed genes expected versus observed in the ILs.

### Differential gene-expression in N2 and CB4856

Differential gene expression between the two parental strains was independently calculated for each environment using a linear model

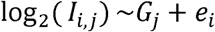

where *I* is the normalized intensity of spot *i* (1, 2, …, 45220) of strain *j* (CB4956 or N2) is explained over genotype *G* and residual variance *e*. For the parental strain N2: n = 4 in control and heat-stress environments, n = 3 in recovery environment. For the parental strain CB4856: n = 3 in control and heat stress environments and n = 5 recovery in recovery environment. The obtained significances were corrected for multiple testing using the Benjamini-Hochberg method implemented in the *p.adjust* function in R (Benjamini and Hochberg 1995). A threshold of FDR = 0.1 was taken as requirement for differential expression.

### Estimation of trait genetic architecture

Two approaches were taken to determine trait architectures using both RIL and IL population: heritability calculations and directly testing the amount of variance between both populations. Both approaches are based on the notion that the trait variance in the populations scale with trait complexity, where

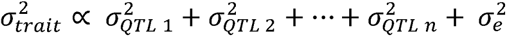

here σ*^2^* is the trait variance, proportional to the variance captured by *n* QTL (which can be multiple loci interacting; epistasis) and the measurement error. The contrast between the RIL and the IL population is that the QTL segregating in the IL population will affect fewer strains per QTL, whereas in the RIL population each QTL will segregate on average in 50% of the strains. In other words, the IL population shares most QTL effects because the genetic background is the same, whereas in the RIL population most QTL segregate.

Heritability was calculated as in (Brem and Kruglyak 2005; Keurentjes *et al.* 2007), where

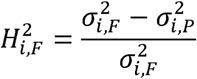

where *H^2^* was the heritability of spot *i* (1, 2, …, 45220) of population *F* (either RIL or IL) within one of the three environments. The σ*^2^* indicated the trait variance of spot *i*. Here the pooled trait variance of the parental lines N2 and CB4856 (denoted with *P*) was used as an estimate of the measurement error, which was subtracted from the trait variance of either the RIL or IL population. We used a permutation approach to determine the threshold for significant heritability (Brem and Kruglyak 2005; Vinuela *et al.* 2012). Thereto, the gene expression values per spot per environment were randomized over the strains (RIL and IL separately, but together with the same parental lines) and recalculated the heritability as described above. This was repeated 1000 times, and the 5% highest value for each spot was taken as the FDR = 0.05 threshold. For the RILs, 4478 spots (2910 genes) displayed significant heritability (FDR = 0.05) in control environment (H^2^ 0.68), 9133 spots (5072 genes) in heat-shock environment (H^2^ 0.70), and 13683 spots (7537 genes) in recovery environment (H^2^ 0.62). For the ILs, 1361 spots (1012 genes) displayed significant heritability (FDR = 0.05) in control environment (H^2^ 0.70), 1647 spots (1123 genes) in heat-shock environment (H^2^ 0.71), and 1578 spots (1092 genes) in recovery environment (H^2^ 0.64).

The second measure was by determining the ratio of trait variance between the IL and the RIL population. The assumption was that the variance contributed by random sources (σ^2^_e_) was equal in both populations. Hence, this is a measure of relative heritability and was calculated as

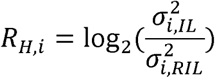

where *R_H_* was the heritability ratio of spot *i* (1, 2, …, 45220), and σ*^2^* was the trait variance in either the IL or the RIL population within one of the tree environments. We took the log_2_ ratio, meaning that traits with a higher heritability in the IL population resulted in a positive number, and traits with a higher heritability in the RIL population with a negative number. We performed two tests to place the relative heritability in perspective. First, we took the spots showing significant heritability (FDR = 0.05). Second, to test whether variance was significantly different between the two populations, we used the non-parametric Fligner-Killeen test for homogeneity of variances, this we corrected for multiple testing using the Benjamini-Hochberg method (FDR = 0.05) (Conover *et al.* 1981; Benjamini and Hochberg 1995).

### Simulating trait architecture impact on relative heritability

To determine how the trait variance behaves proportionally in each population, we simulated different trait architectures in both the IL and RIL population. First, all QTL effect sizes were simulated by random drawing from a standard normal distribution (µ = 0, σ = 1). These QTL were simulated in trait architectures containing 1 to 100 QTL (or 1 to 100 clusters of closely linked QTL, each 33331 times. To recapitulate the observed QTL distribution in *C. elegans*, we distributed these QTL along the informative markers in our IL and RIL population. This assured an enrichment of QTL on the chromosome arms compared to the chromosome tips and centers (Rockman *et al.* 2010; Snoek *et al.* 2017). To approach the residual variance 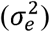 in our data, we added random variation based on a normal distribution (*rnorm*) such that the average heritability of each simulated trait was 0.85

The following eight architectures were simulated: (i) randomly distributed additive QTL, (ii) randomly distributed additive QTL that only show their effect in one genotype, (iii) clusters of closely linked QTL with random effects, (iv) clusters of closely linked QTL that only show their effect in one genotype, (v) clusters of closely linked QTL that are balanced within the ancestral genotypes, (vi) clusters of closely linked QTL that are balanced within the ancestral genotypes and only show their effect in one genotype, (vii) randomly distributed interacting QTL, (viii) clusters of closely linked interacting QTL. The number of closely linked QTL per cluster was simulated from two to five. The additive and closely-linked model were similar in essence, but differ in the architecture, where we modelled the closely-linked QTL such that these were typically not segregated in the population used.

The distribution of traits over a balanced closely-linked architecture and additive architecture was done by measuring the number of heritable traits with a heritability ratio between −3 and −2 (peak) and between 0.5 and 1.5 (intersect). These intervals were chosen as they represent areas where the distributions of these models differ. The number of expected traits per interval were counted as the median of all simulated architectures (so 1 to 100 QTL, and 2 to 5 QTL per closely-linked locus) in order to get an estimation that is less dependent on the chosen simulation parameters. The predicted number of traits at the peak and the intersect for the additive model (a) and the balanced closely-linked model (b) were used to solve the true distribution based on the real number of heritable traits in these heritability ratio intervals (c). This was done by solving the equations

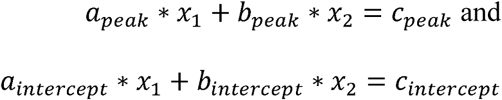

for x_1_ and x_2_ using the *solve* function in R. This led to the estimated number of additive traits (x_1_) and balanced closely linked traits (x_2_).

## Results

### Genome-wide confirmation of eQTL effects and location using introgression lines

Analysis of differential expression showed that introgression lines (ILs) could confirm both *cis*- and *trans*-eQTL mapped in the recombinant inbred line (RIL) population (Snoek *et al.* 2017). A population of 48 RILs, 56 ILs ((Doroszuk *et al.* 2009); **Table S1**), and at least three replicates of the ancestral strains N2 and CB4856 were exposed to the three environments (Figure 1A). Based on the number of eQTL expected in the CB4856 locus of each IL, we found that on average 69.5% (FDR = 0.05) of the *cis*-eQTL found using the RILs could be confirmed using the ILs over all three environments (FDR = 0.05). The *trans*-eQTL found in the RILs, were confirmed on average 42.2% (FDR = 0.05) in the ILs (Figure 1A and 1B; **Table S2**). Note that the tiling structure of the ILs meant that each QTL location was covered by 0-5 (5-95% quantiles) ILs (**Figure S1**). The percentage of confirmed eQTL was positively correlated with the RIL significance threshold (**Figure S2A**). Furthermore, correlation analysis of the log_2_ ratio with the N2 ancestor in the ILs versus the eQTL effect measured by RILs showed that eQTL effects in the CB4856 locus of each strain were well approximated in the ILs (average Pearson correlation, rho = 0.661; **Figure S2B**; **Table S2**). Together, over all three environments, the analyses confirmed eQTL occurrence and effect sizes in the RIL population in independent experiments using an IL population.

**Figure 1:**
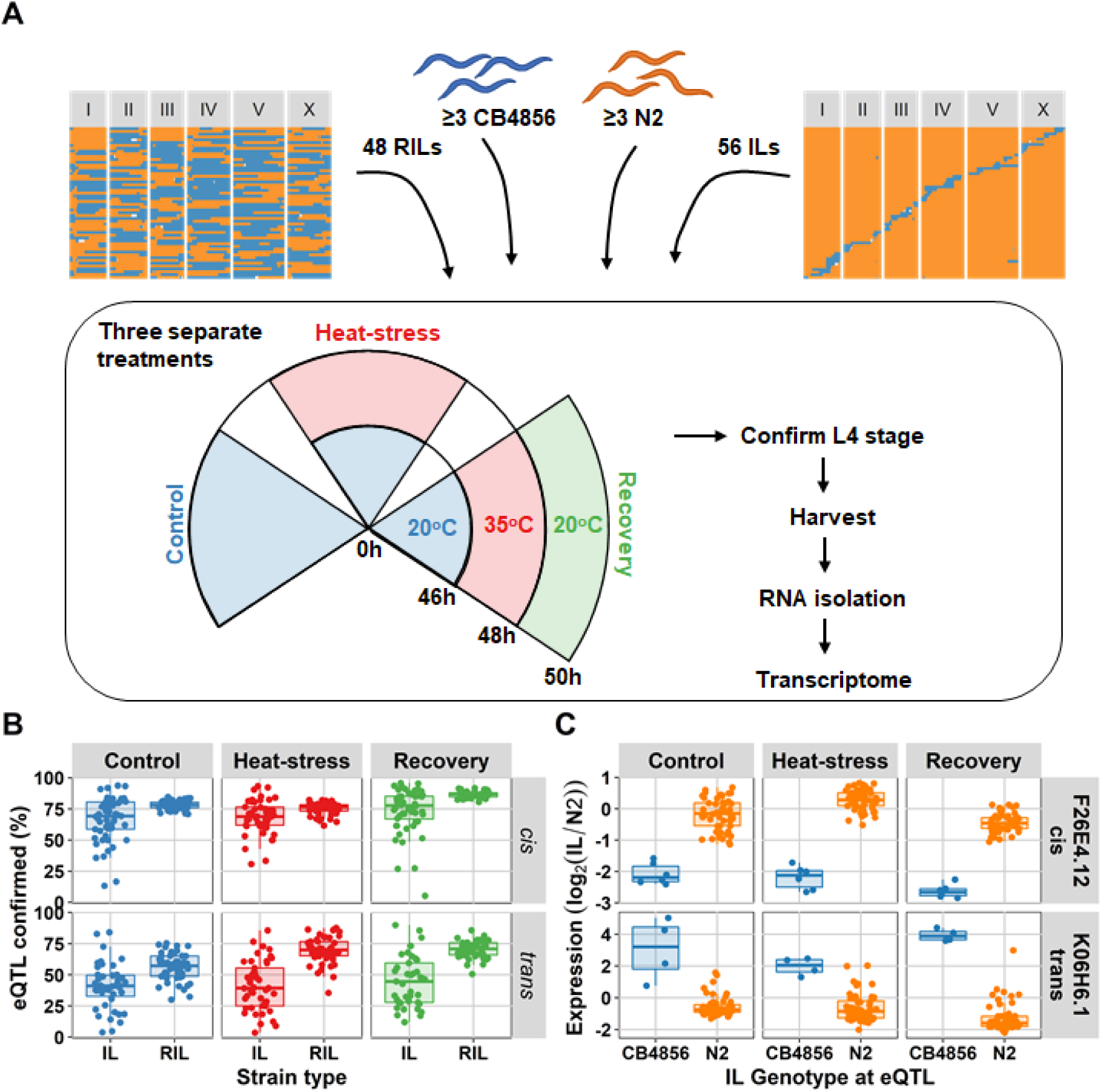
Experimental setup and eQTL confirmation (**A**) The four types of populations used in the experiment: recombinant inbred lines (RILs), the ancestral strain CB4856, the ancestral strain N2, and introgression lines (ILs). These were exposed to three separate treatments: control (48 hours at 20°C), heat-stress (46 hours at 20°C followed by 2 hours at 35°C), and recovery (as heat-stress, followed by an additional 2 hours at 20°C). After the experiment, populations of the strains were harvested and used for transcriptome measurement by microarray. (**B**) the percentage of eQTL confirmed based on differentially expressed genes per IL and per RIL (FDR = 0.05). The percentages are split-out over the three environments, and the two eQTL type (*cis* or *trans*). The eQTL were mapped using the RIL population, hence that population is expected to confirm the eQTL to a high degree. (**B**) An example of one *cis*-eQTL (for gene F26E4.1) and one *trans*-eQTL (for gene K06H6.1) in the IL population. On the x-axis the genotype at the QTL locus is plotted, and on the y-axis the expression (R_N2_)

Next, we investigated the *trans*-bands identified in the RIL population more closely. The *trans*-bands were described in a previous study (Snoek *et al.* 2017). For each *trans*-band, we took 2 to 6 ILs covering and flanking the locus and tested if the *trans*-eQTL effects correlated significantly with differentially expressed genes in the ILs compared to N2. We found supporting evidence for 14 out of 19 *trans*-bands at an overall false-positive rate of 0.11; see materials and methods. Furthermore, we narrowed down the location for some of these *trans*-bands (**File S1**; **Table S3**). In particular the *trans*-bands in the control and heat-stress environment were well supported (5/5 and 6/7, respectively), whereas those in the recovery environment were less (3/7). The poor support in the recovery environment is likely due to the few *trans*-eQTL per *trans*-band in that environment (median of 25 genes in recovery versus 51 in control and 125 in heat stress). Altogether, we conclude that *trans*- bands were highly replicable, to a higher extent than individual *trans*-eQTL.

Next, we asked if the number of differentially expressed genes in the IL population could be predicted from the RIL population. Therefore we compared the gene expression in both populations to the CB4856 ancestor to measure the effect of the N2 loci and to the N2 ancestor to measure the effect of the CB4856 loci. The effect of CB4856 introgressions in an N2 genetic background on gene expression was negatively correlated with the expectations based on the RIL population. In other words, in the RIL population the number of genes differentially expressed due to the introgression was under-estimated and the number of genes differentially expressed due to the genetic background was over-estimated (Figure 2). In the ILs we found that the relative number of differentially expressed genes compared to N2 was 6.7-fold higher than expected based on the differentially expressed genes found in the RILs (paired t-test, p < 1*10^-54^). Furthermore, when compared to CB4856, the number of differentially expressed genes in the ILs was 3.3-fold lower than expected from the RILs (paired t-test, p < 1*10^-128^). The differences between RILs and ILs were consistent when changing both the threshold for differential expression and the threshold for eQTL-based expectations (**Figure S3**). These results show that the expectations from a single marker model are a poor predictor for differentially expressed genes in ILs. Importantly, a single marker model under-estimates the impact of small introgressions and over-estimates the impact of the genetic background.

**Figure 2:**
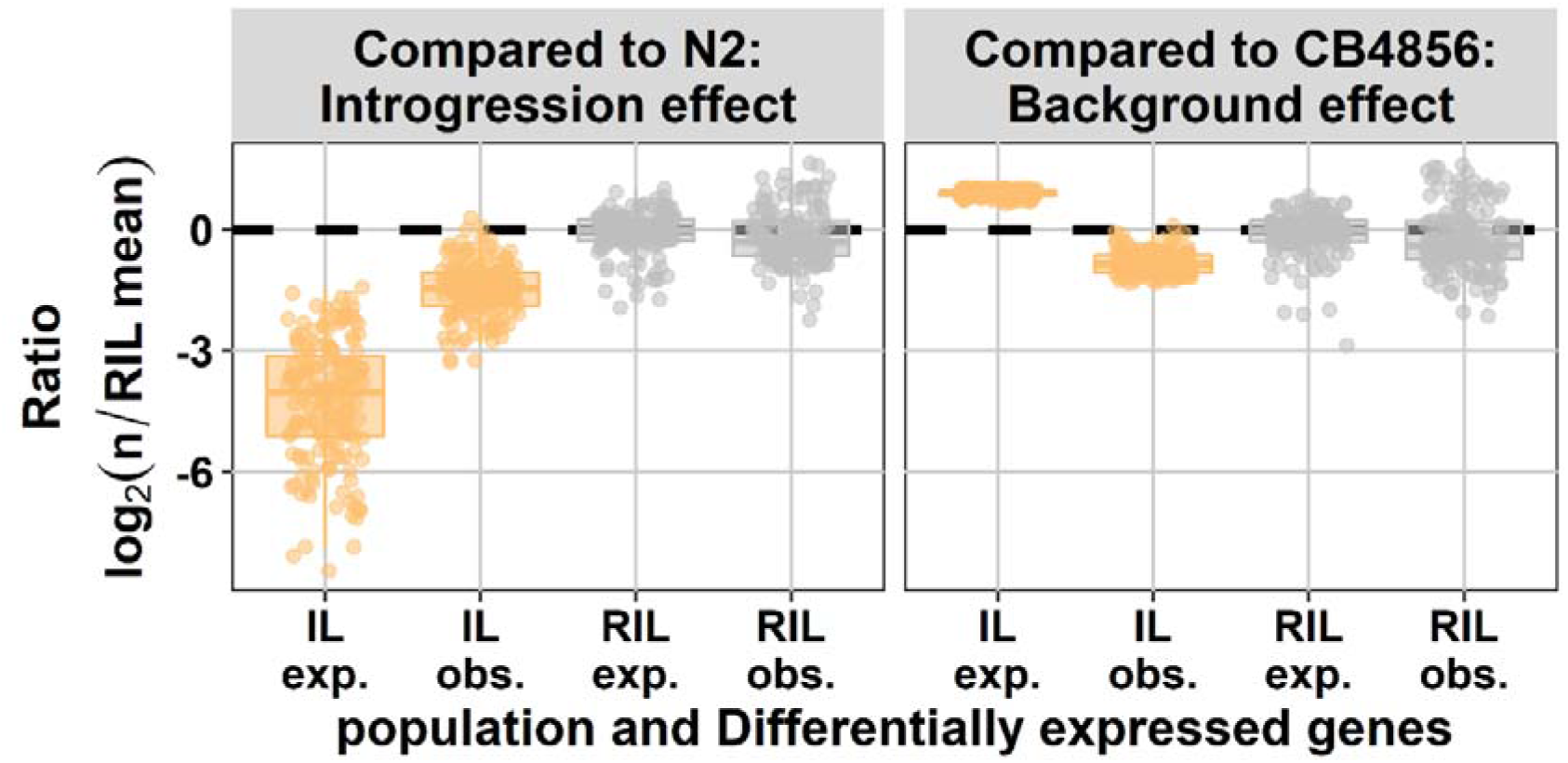
Number of differentially expressed genes (DEG) expected (exp.) from eQTL mapping and observed experimentally (obs.). On the x-axis, the population type is shown and on the y-axis, the expected and observed ratio of differentially expressed genes is shown, both normalized for the expectation in the RILs (eQTL threshold, -log_10_(p) > 3.9; DEG threshold, - log_10_(p) > 5). In the left panel the effect compared to N2 is shown (effect of the CB4856 loci; introgressions in the IL panel) and in the right panel the effect compared to CB4856 is shown (effect of the N2 loci; genetic background in the IL panel).

### Micro-architecture of eQTL

The distribution and differences of genetic recombination over ILs and RILs allowed studying the architecture of eQTL in *C. elegans*. We hypothesized that different underlying trait architectures would leave different trait-variance distributions in the IL and RIL population. For example, a simple trait architecture dominated by one major QTL would only lead to an effect on trait levels in a single or a few ILs, but in ∼50% of the RILs. Hence, resulting in a low trait variance over the genome-wide IL population compared to the RIL population. As heritability was a straightforward way to interpret trait variance, we estimated the broad-sense heritability (H^2^) in both populations (**Table S4**). In general, heritability was higher in the RIL population. In total, over all conditions, we found 10673 unique genes that showed significant heritability in the RIL population, whereas we found 2952 unique genes that showed significant heritability in the IL population (permutation, FDR = 0.05; **Figure S4A**). Only 929 genes in the RILs and 8 genes in the ILs showed heritable gene expression variation over all three environments (**Figure S4B** and **C**). The reason for the difference between populations was partly due to 534 out of 929 genes having an eQTL in the RILs (of which 354 had a *cis*- eQTL), which captured little heritable variation in the whole IL population. From the heritability analysis, we conclude that, as expected, heritability was higher in the RIL population, and that heritability is highly environment dependent.

To gain insight in the impact of different trait architectures, we simulated eight distinct architectures, including additive and epistatic architectures (see Materials and methods, **Figure S5**, and **Figure S6**) and measured trait variance ratios between the IL and RIL population. Subsequently, we assumed that the measurement error 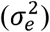 was similar for both populations and calculated the heritability ratio between both populations (Figure 3A). Based on the additive simulation, we expected 95% of the variance ratios to fall between –3.6 and – 1.7, whereas under a model with closely-linked QTL that cancel each other out in the same genetic background (balanced), the 95% interval falls between –24.9 and 4.9. The balanced closely-linked QTL architecture was modelled such that QTL within such a cluster would only separate occasionally by a recombination in the RIL and IL population. The shape of the distribution of genes with significant heritability seems to form a combination of additive and balanced closely linked QTL (Figure 3A). To get an estimate of the contribution of the two models, we used the difference in the additive and balanced closely linked distributions from the simulation to estimate how the traits are divided over the two QTL models (Figure 3B). Analysis of the two distributions revealed that in all three environments additivity was the most likely QTL model (60 – 93% of genes with significant H^2^). Balanced closely linked QTL explained up to 40% of relative heritability in the control environment, but only 7% in the heat stress and recovery environments (**Table S5**). Therefore, we hypothesize that the balanced closely linked QTL model is an important eQTL architecture in the nematode *C. elegans*.

**Figure 3:**
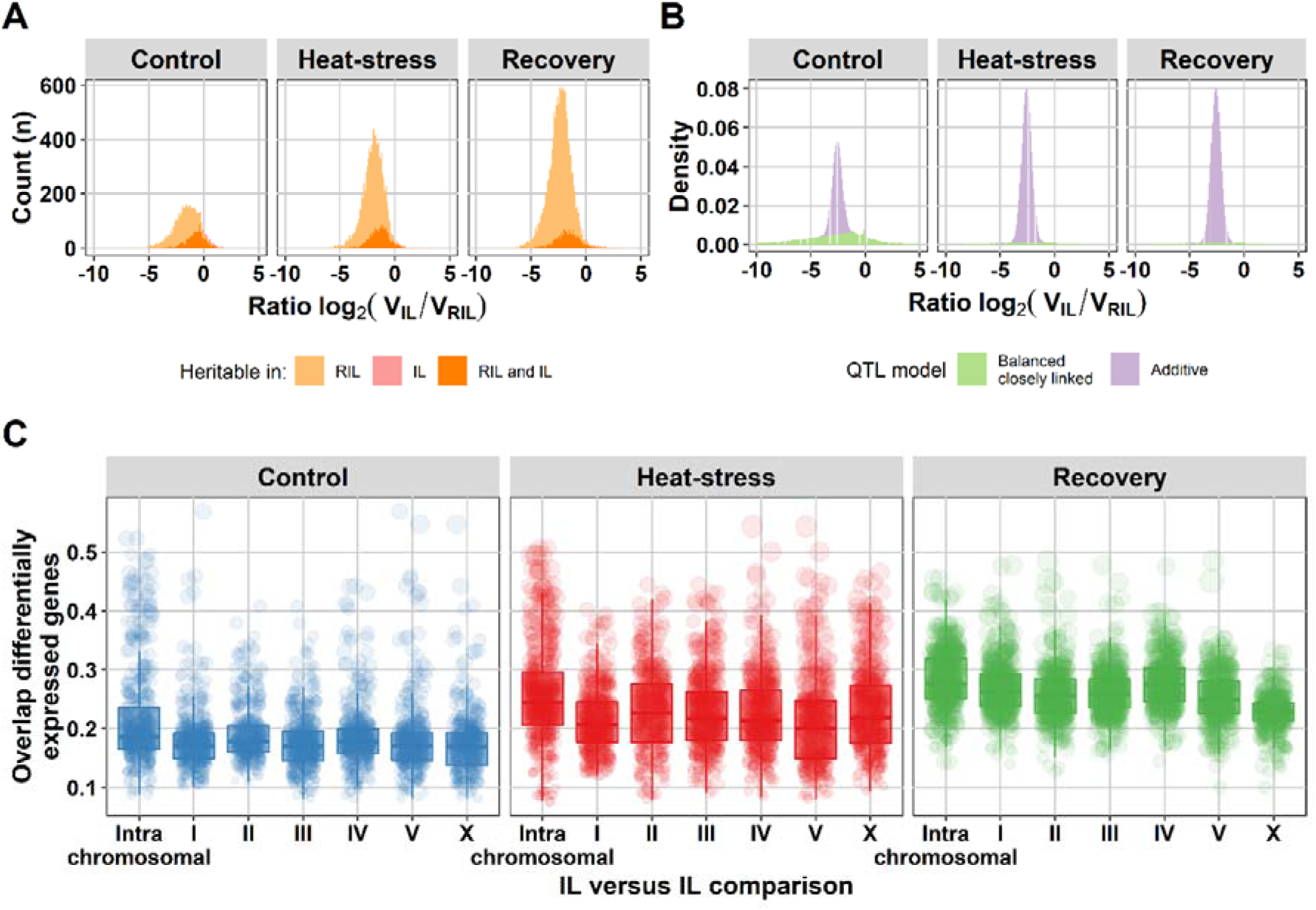
Balanced closely linked QTL can explain heritability ratios between the RIL and IL populations. (**A**) Histogram of the heritability ratio in the IL versus the RIL population of genes with significant gene expression heritability (permutation, FDR = 0.05). The light orange surface indicates microarray spots of genes with heritable variation in gene expression only in the RIL population (3262 in control, 7545 in heat-stress, and 12176 in recovery), the pink surface microarray spots of genes with only heritable variation in gene expression in the IL population (145, 59, and 71 respectively) and the orange color microarray spots of genes with heritable variation in gene expression in both populations (1216, 1588, and 1507 respectively). (**B**) The density distribution of the heritability ratios of genes with heritable gene gene expression variation over the balanced closely linked (green) and the additive (purple) QTL model. Trait models consist only of additive or balanced closely linked QTL. (**C**) Overlap in differentially expressed genes between ILs (threshold –log_10_(p) > 5). The comparisons are grouped per chromosome (x-axis): the intra-chromosomal comparisons and all other chromosomes versus chromosome I, II, III, IV, V, or X. On the y-axis the fraction overlap is shown. The dots represent one IL versus IL comparison the size corresponds to the number of overlapping differentially expressed genes.

### Two predictions from the balanced closely linked eQTL microarchitecture

The balanced closely linked QTL model comes with two testable predictions: (i) the genes with heritable variation in gene expression were not differentially expressed in the parental strains, (ii) differential expression of genes was not only linked to one location in ILs but to clusters of ILs perturbing distant loci (multiple balanced clusters). Indeed, only 9.1 to 15.4% of the genes with heritable variation in gene expression were differentially expressed when comparing the two parental strains (FDR = 0.1; **Table S6**). Also, the second prediction was found to be correct, on average 21.9% of the differentially expressed genes were shared between IL strains covering loci on different chromosomes, which was only slightly less than the average of 25.1% between ILs covering the same chromosome (Figure 3C and **Figure S7**, and for examples Figure 4). Therefore, we concluded that localized genetic complexity was a major part of the eQTL architecture.

**Figure 4:**
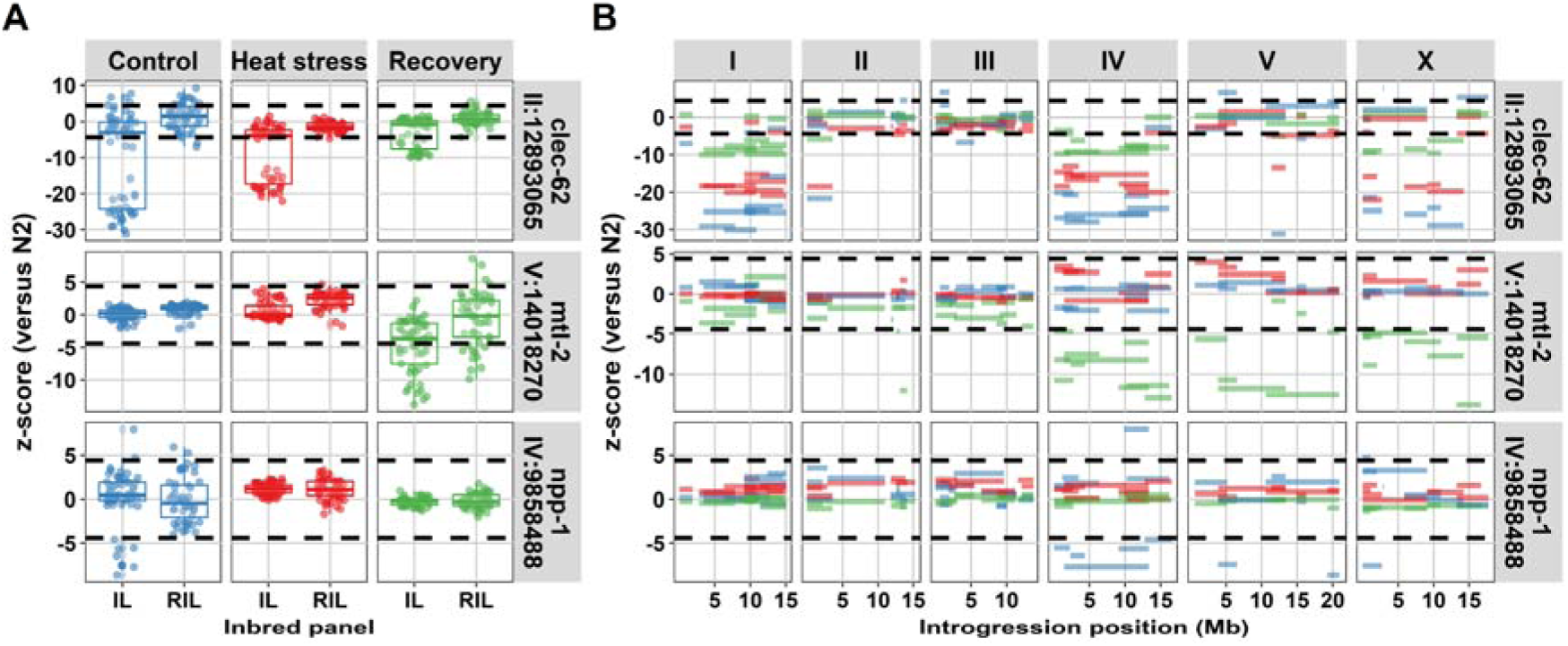
Three genes with high heritability in gene expression variation in the IL population compared to the RIL population. (**A**) Comparison in expression between IL and RIL population for *clec-62*, *mtl-2*, and *npp-1*. On the x-axis, the population type and on the y-axis, the z-score with the N2 parental lines. The dashed horizontal lines indicate the significance threshold (z-score, -log_10_(p) > 5). (**B**) The expression per introgression. On the x-axis, the introgression location is shown per chromosome. On the y-axis, the z-score with the N2 parental line is shown. The dashed horizontal lines indicate the significance threshold (z-score, -log_10_(p) > 5). The colors indicate the environments (blue for control, red for heat-stress, and green for recovery).

Furthermore, differences in heritability between the RIL and IL populations over environments indicated that the control environment was different from heat-shock and recovery. The balanced closely linked architecture, which predicted a broad range of relative heritability, seemed to be most prominent in the control environment. It could be that these complex architectures were more important for the developmental process. This environmental effect could be illustrated by the expression variation of the genes: *mtl-2*, *npp-1*, and *clec-62*. Of which *mtl-2* and *npp-1* showed an interaction with the environment. The gene *npp-1* only showed this interaction in the control environments, environment where ILs on multiple chromosomes showed differential expression of this gene, yet for *mtl-2* ILs with differential expression on different chromosomes were found during recovery conditions, whereas *clec-62* shows ILs with differential expression on different chromosomes in control and heat stress conditions. (Figure 4). The nuclear core complex protein NPP-1 is involved in spindle formation and important for oogenesis, a process that was interrupted during heat-shock (Schetter *et al.* 2006; Jovic *et al.* 2017). MTL-2, one of two metallothioneins in *C. elegans*, was expressed in the intestine and was induced after heat-shock (Freedman *et al.* 1993). The pattern observed for *clec-62* was exceptional as it showed a consistent response over all three environments where IL strains with an introgression on chromosome I, chromosome IV, and chromosome X showed much lower expression compared to the N2 ancestor. Furthermore, also an IL strain with an introgression on chromosome II and one with an introgression on chromosome V displayed this phenotype. C-type lectin 62 is a gene belonging to the extensive *clec*-family, which is thought to be involved in *C. elegans* immunity (Schulenburg *et al.* 2008) although its exact function is not known.

## Discussion

### Most eQTL mapped in RILs are replicable in ILs

Here, we show that eQTL mapped in a RIL population can be confirmed in a different type of inbred line population. We found that the majority of *cis*-eQTL are replicable in the IL population (on average 69.5% per IL), whereas *trans*-eQTL are less replicable in ILs (on average 42.2% per IL). This is likely to reflect the mainly monogenic architecture of *cis*- eQTL, versus the more polygenic and environment-dependent *trans*-eQTL (Li *et al.* 2006; Keurentjes *et al.* 2007; Rockman *et al.* 2010; Snoek *et al.* 2017; Albert *et al.* 2018). Especially the confirmation of *trans*-bands by ILs shows that these regulatory hot-spots are robust regardless of genetic background. It should be noted that this study does have limited power per introgression line as each line was only measured once per condition. Hence we avoid conclusions based on a single ILs throughout this paper. However, the tiling nature of the population does mean that each locus is covered by multiple ILs (Doroszuk *et al.* 2009).

This study helps the search for candidate genes underlying these *trans*-bands, by narrowing-down the regions. However, additional experiments and replicates are recommendable and currently pursued for the heat-stress *trans*-band on chromosome IV. Furthermore, it shows how ILs can be used to narrow-down these eQTL hotspots (Snoek *et al.* 2012), using correlation analysis previously used to link *trans*-bands to genes and biological processes (Andersen *et al.* 2014; Sterken *et al.* 2017). These findings show on a large scale that QTL mapped using a single marker model in a moderately sized RIL population are reliably replicable in a population with a different genetic structure, which confirms findings for single traits reported in *C. elegans* and beyond (for example, see (Snoek *et al.* 2012; Andersen *et al.* 2014; Gao *et al.* 2018)).

### Introgression lines reveal the presence of parental-balanced, polygenic, traits

The difference in genetic complexity between RILs and ILs can be leveraged to understand trait architectures. Despite confirming many QTL, it has been noted that ILs often tell a different story than RILs, implying abundant genetic interactions, not uncovered by RILs (as reviewed by (Mackay 2014)). In this study, we also show that the number of differentially expressed genes in the N2 genetic background of the ILs is lower than expected compared to RIL-based estimations. Furthermore, eQTL mapped in the RIL population also led to an underestimation of the number of differentially expressed genes due to the introgressions in the ILs. To summarize, the introgressions display more-than-additive effects, whereas the genetic background shows less-than-additive effects.

Our findings are in line with observations in introgression lines in *C. elegans*, showing that for some traits more and different QTL than expected from RILs can be found (*e.g.* (Gaertner *et al.* 2012; Glater *et al.* 2014; Snoek *et al.* 2014a)). Additionally, our findings are also in line with findings in other organisms, where such effects have been reported for different types of traits (Shao *et al.* 2008; Gale *et al.* 2009; Spiezio *et al.* 2012). In this study, we show this effect in general, over many gene expression traits. Moreover, a direct comparison of trait mapping in a genome-wide IL population versus a RIL population has only been conducted in a few studies, as far as we are aware (Glater *et al.* 2014; Snoek *et al.* 2014a). It has been argued that ILs showing more- or less-than-additive effects compared to the parental strains is a hallmark of epistasis (as reviewed by (Mackay 2014)). However, recent studies in yeast show that additivity underlies most of the heritable trait variation among inbred lines, where epistasis accounts for phenotypic extremes (Bloom *et al.* 2015; Forsberg *et al.* 2017; Albert *et al.* 2018). Thus remains the question, what kind of trait architecture drives our observations: pervasive epistasis or widespread additivity?

### The developmental trait architecture consists of balanced closely linked QTL

Trait architectures underlying natural variation in *C. elegans* differ strongly over traits. For example, currently 25 quantitative trait nucleotides (QTNs) are known in *C. elegans*, capturing a majority of the heritable variation for particular traits (reviewed by (Rockman 2012) and (Gaertner and Phillips 2010) and studies by (Andersen *et al.* 2014; Noble *et al.* 2015; Schmid *et al.* 2015; Cook *et al.* 2016; Greene *et al.* 2016a; Greene *et al.* 2016b; Large *et al.* 2016; Ben-David *et al.* 2017; Sterken *et al.* 2017; Zdraljevic *et al.* 2017; Hahnel *et al.* 2018; Zdraljevic *et al.* 2019). However, there are also examples of traits that are highly heritable but have only yielded complex or few QTL (let alone QTNs). For example, a study on bacterial preference of *C. elegans* noted a relatively high heritability (0.46) for *Serratia marcescens* over *E. coli*, yet uncovered only a single QTL in the RILs used for mapping whereas multiple were expected (Glater *et al.* 2014). Furthermore, recent work from our group, studying metabolite abundances showed high heritability (from 0.32 up to 0.82) corresponding to a few uncovered QTL (Gao *et al.* 2018). Intriguingly, several studies imply trait architectures consisting of closely linked QTL that are balanced in the parental strains (Gaertner *et al.* 2012; Glater *et al.* 2014). By simulation, we show that some trait architectures can be differentiated by relative heritability between IL and RIL populations which is a property we can reliably measure in our two populations, as it only relies on determining trait variance. In this way, we found that for genes without an eQTL, but showing high heritability, the most parsimonious explanation lies in an architecture comprised of (multiple) closely linked QTL clusters. This type of architecture seems especially prominent during normal development where it could affect approximately 40% of genes with heritable expression variation. It should be noted that only strictly balanced closely linked and strictly additive QTL distributions were modelled. We think the observed distribution suggests that a combination of (both or multiple) types of QTL make up the complete genetic architecture of a trait.

In recent years, the number of parameters known to affect gene expression variation in the context of natural variation has steadily grown. It is currently clear that environment (Li *et al.* 2006; Snoek *et al.* 2017), age (Vinuela *et al.* 2010), development (Francesconi and Lehner 2014), and background mutations (Sterken *et al.* 2017) affect gene expression variation and eQTL distribution in *C. elegans*. These factors could also contribute to the observed heritability differences between ILs and RILs. For environment, we can confirm that it plays an important role: the heritability ratio is strongly dependent on environment. This is in line with effects observed for *trans*-eQTL, which are also strongly environment dependent, implying heritability is as well (Li *et al.* 2006; Snoek *et al.* 2017). Developmental differences between the strains in each population could also drive some of the observed heritability ratios. For example, ILs could be developmentally more homogenous than RILs, which could result in lower levels of trait variance in ILs. It is clear that age affects *trans*-eQTL and heritability (Vinuela *et al.* 2010; Vinuela *et al.* 2012). However, it is unclear how that would affect the heritability ratios we measure in general; it seems unlikely that developmental differences result in a specific variance signature as changes occur in both populations. More specific, in this experiment strains are tightly synchronized and developmental stage is observed to be L4 before isolation, therefore we expect that these potential sources of increased variance are controlled. Hence, the relative heritabilities under normal development imply clusters of closely linked QTL.

### The role of cryptic genetic variation on trait complexity

We used the contrast in genetic complexity of the RIL and IL population to investigate trait complexity. The relative heritabilities between these populations differed over the three environments. In our previous study on the RILs, we show that the *trans*-eQTL architecture is different over the three environments, showing that *trans*-eQTL contribute most to cryptic genetic variation. Here, we find that the micro-architectures is more complex in an ambient environment.

Development of an organism is a tightly regulated process and completion of development into a reproducing adult is essential for reproduction, hence fitness. The nematode *C. elegans* experiences outbreeding depression (Dolgin *et al.* 2007), given the strains N2 and CB4856 are genetically distinct, it is likely that inbred lines constructed from these two strains disrupt some linked loci involved in this process (Seidel *et al.* 2008; Snoek *et al.* 2014a). We hypothesize that especially the ambient environment stimulating normal development is prone to accentuate the effects of genetic ‘mismatches’ resulting in trait variation. First, local sites are expected to co-segregate over many generations in *C. elegans*, making it likely that they are filled with compensatory mutations (Rockman *et al.* 2010). Second, development is a polygenic, tightly regulated, process. On the level of gene expression, there are strong changes over the entire development from egg to adult, and during the last juvenile stage (L4) especially (Francesconi and Lehner 2014; Hendriks *et al.* 2014; Snoek *et al.* 2014b). Together, these two effects may cause the *C. elegans* populations in ambient environments to show micro-architectures that are more complex.

On the other hand, stress responses are also tightly regulated, but initially aimed at reaching a stress-survival mode, which is a process that involves fewer genes than development. The heat-shock response in *C. elegans* starts by de-phosphorylation of HSF-1 and DAF-16 and entry of these transcription factors into the nucleus (as reviewed by (Rodriguez *et al.* 2013)). Once there, they regulate the expression of chaperones that function to minimize cellular damage. Our results show that populations in both the heat-shock and recovery from heat-shock environments show fewer QTL with complex micro-architectures. We hypothesize that this is due to the effect of the strong transcriptional response induced by heat shock (Brunquell *et al.* 2016; Jovic *et al.* 2017). It is possible that this abrupt disturbance emphasizes regulatory variation in only a few key response pathways compared to the more subtle and complex process of development.

## Conclusions

We present an eQTL experiment conducted with ILs in three environments covering the same conditions as a RIL experiment published previously (Snoek *et al.* 2017). We show that ILs can replicate both *cis-* and *trans*-eQTL; furthermore, we present evidence supporting 14 eQTL hot spots (*trans*-bands). Yet, eQTL mapped in the RIL population systematically under-estimate the impact of a single introgression on the transcriptome and over-estimate the impact of the genetic background, suggesting additional genetic complexity. We show that during normal growth multiple clusters of closely linked QTL are underlying additional genetic complexity.

Further understanding of this phenomenon requires systematic dissection of many QTL – a process that in this study was only undertaken for *trans*-bands – using introgression lines. It remains to be determined what kind of role epistasis plays in trait variation. Currently, RIL populations in most species are typically of insufficient size to detect any epistasis beyond two loci interactions. Careful crosses with ILs to generate double-loci ILs might be a way forward to further our understanding of trait regulation in the context of natural genetic variation.

## Acknowledgements

The authors thank the GRAPPLE project partners: Olga Valba, Sreenivas Chavali, Benjamin Lang, Mirko Francesconi, Rachel Brenchley, Arjen van’t Hof, Sergei Nechaev, Olga Vasieva, Madan Babu, Andrew Cossins, and Ben Lehner. We thank Harm Nijveen for assistance with WormQTL2.

## Funding

LBS was funded by ERASysbio-plus ZonMW project GRAPPLE - Iterative modelling of gene regulatory interactions underlying stress, disease and ageing in *C. elegans* (project 90.201.066) and The Netherlands Organization for Scientific Research (project no. 823.01.001). JEK was funded by NIH grant 1R01AA 026658-01. The funding bodies had no role in the design nor the collection, analysis, and interpretation of the data, nor the writing of the manuscript.

## Availability of data and materials

The strains used in this study can be requested from the authors. The transcriptome datasets used in the analysis are deposited at ArrayExpress E-MTAB-5779 and E-MTAB-7424.

## Author contributions

MGS, LBS and JEK conceived and designed the experiments. MGS wrote the manuscript, with input from JEK and LBS. MGS, RPJB, RJMV, and JAGR performed the experiments. MGS analyzed the data, with input from LBS. All authors commented on the manuscript.

## Supplementary information

**Figure S1:** Coverage per locus and per QTL. (**A**) the coverage in CB4856 loci per location on the genome, split out for ILs and RILs. The 56 ILs together have a higher coverage over the chromosome arms, where also most QTL map. The 48 RILs have a more homogenous distribution, only at the *peel-1*/*zeel-1* locus on chromosome I there is low coverage (Seidel *et al.* 2008). (**B**). A histogram of the number of CB4856 loci covering an eQTL. Typically, an eQTL is covered by CB4856 loci of 2 ILs and 23 RILs (median).

**Figure S2:** Confirmation of eQTL by significance thresholds and correlation analysis. (**A**) The percentage of confirmed eQTL per population per eQTL-type. On the x-axis, the -log_10_(p) significance threshold in the RILs is shown (binned per 0.5 in the analysis) and on the y-axis the percentage of confirmed eQTL per bin per strain is shown (at least five confirmable eQTL required per bin). For example, for the RIL WN98 under recovery conditions there were 115 *cis*-eQTL with a significance between 4.0 and 4.5 in its CB4856 regions, 91 of these were confirmed based on the z-score (79.1%). This example results in a dot where the size corresponds to the number of eQTL and the color indicates the environments. The line shown is a linear regression curve, where the surrounding grey area is the confidence interval. (**B**) The Pearson correlation of eQTL effects with gene expression in the ILs and the RILs. On the x-axis the population type is plotted and on the y-axis the Pearson correlation. The correlations are split-out over the three environments (control, heat stress, and recovery) and over eQTL type (*cis* or *trans*).

**Figure S3:** Differentially expressed genes in RILs and ILs. (**A**) Number of differentially expressed genes (DEG) expected from eQTL mapping and observed experimentally, split out per environment. On the x-axis the population and on the y-axis the expected and observed ratio of differentially expressed genes is shown, both normalized for the expectation in the RILs (eQTL threshold, -log_10_(p) > 3.9; DEG threshold, -log_10_(p) > 5). In the top panel the effect compared to N2 is shown (effect of the CB4856 loci; introgressions in the IL panel) and in the bottom panel the effect compared to CB4856 is shown (effect of the N2 loci; genetic background in the IL panel). (**B**) The effect of different thresholds for eQTL significance (x-axis) and DEG calling (y-axis) on the ratio between observed and expected.

**Figure S4:** Genes with heritable variation in gene expression in the RIL and IL populations. (**A**) A histogram of the measured heritabilities for the IL and RIL populations, negative values have been discarded. Colours indicate whether the heritability was significant (FDR = 0.05) in the control (blue), heat-stress (red), or recovery (green) environment. (**B**) The overlap in genes with significant heritability in gene expression variation over the environments for the RIL population (FDR = 0.05). (**C**) The overlap in genes with significant heritable variation in gene expression over the environments for the IL population (FDR = 0.05).

**Figure S5:** A graphical overview of the modelled trait architectures. The arrows indicate QTL effect sizes as found in the N2 genotype (orange) and the CB4856 genotype (blue). Perfectly opposed arrows, one of which orange and the other blue, indicate that the same QTL has an opposite effect in the two genotypes (e.g. as seen for Additive random distribution). When an arrow lacks a perfectly opposed arrow of another colour, it means that the QTL is only found in that particular genotype. Epistatic interactions are indicated as a line connecting two rectangles.

**Figure S6:** Variance ratios between IL and RIL population for the simulated trait architectures (**Figure S5**). The density of occurrence of variance ratios per simulation is given. The colour scale indicates how many QTL for a trait were simulated.

**Figure S7:** Overlap in differentially expressed genes between ILs (threshold –log_10_(p) > 5. The ILs are ordered based on introgression on the x- and y-axis. The location of the CB4856 segment is indicated behind the IL strain-code. The fraction overlap shown is calculated as the percentage of unique differentially expressed genes in the two compared ILs. The overlap is shown for the control environment (**A**), the heat-stress environment (**B**), and the recovery environment (**C**).

**File S1**: Comparison of gene expression in ILs with eQTL in *trans*-bands identified in the RIL population. Per *trans*-band, a figure was constructed. In (**A**) the name of the *trans*-band is given at the top of the panel (e.g. chromosome I, 3.5-4 million bases) the correlation of the relative gene expression (R_N2_) was correlated with the eQTL effect for the ILs covering or nearby the *trans*-band. Significance was determined based on the highest correlation in the N2 strains, only strains with a stronger correlation than N2 were scored as significant. In (**B**) the genetic map of the ILs of the *trans*-band chromosome is shown. On the x-axis the genomic location (in million bases) is shown and on the y-axis the ILs. In (**C**) the actual correlation between expression in the ILs and eQTL effect in the RILs is shown. With on the x-axis the eQTL effect and on the y-axis R_N2_. Each dot represents a spot on the microarray with an eQTL in the *trans*-band investigated. The line indicates the estimated slope and the pale blue area around the line indicates the confidence interval of the fit.

**Table S1:** The genotypes of the strains used in this study, −1 denotes a CB4856 genotype, a 1 denotes an N2 genotype. The genotypes are given per chromosome and per position (based on WormBase version WS258)

**Table S2:** Table of the eQTL with gene expression comparisons per strain per environment. The table lists the strain name, strain type and the comparisons are split out for the *cis*- and *trans*-eQTL. The correlation reported is the Pearson correlation and its significance. The number of eQTL present in the CB4856 loci is given, as is the percentage that is also differentially expressed (FDR = 0.05).

**Table S3:** Summary of the comparison of gene expression in the ILs with the *trans*-bands identified previously (Snoek *et al.* 2017). The location of the *trans*-band and the number of genes and microarray spots affected are listed. The genotypes of the ILs with an introgression adjacent to – or covering the *trans*-band are listed. The genotypes of the ILs that confirm the *trans*-band and the *trans*-band locus that these ILs imply are also given.

**Table S4:** Table with the heritabilities per spot per environment per population. The estimated genetic variance (V_g_) and the technical variance (measurement error, V_e_) are given, as is the heritability (H^2^). The FDR0.05 column indicates the threshold of significance based on permutation.

**Table S5:** A table with the estimated fraction and amount of traits for an additive QTL architecture and a balanced closely-linked QTL architecture. The numbers are calculated based on only a single type of QTL occurring for a particular architecture. For example, additive means *n* QTL that display an additive effect, without any other type of effect for that simulated trait.

**Table S6:** differentially expressed genes between N2 and CB4856 for the three environments: control, heat-stress, and recovery from heat stress. The columns in the table: SpotID, the Agilent spot identifier, the information on the gene expression detected by that spot is given (chromosome, location (start/stop), strand, and three identifiers). Two columns show if genes are significantly heritable; general (in any population) and specific (which population, or both). The significance and effect columns give the output of the linear model when comparing CB4856 versus N2 (positive effect indicates the gene is higher expressed in N2). The last column gives the significance as corrected for multiple testing (FDR).

## Notes

#### Summary of Updates

The text has been expanded to clarify analyses, a false-discovery rate was calculated for the trans-band verification.

https://git.wur.nl/published_papers/sterken_2019_closely_linked_qtl

https://www.ebi.ac.uk/arrayexpress/experiments/E-MTAB-5779

https://www.ebi.ac.uk/arrayexpress/experiments/E-MTAB-7424

