## Supplementary material for "Dissecting the eQTL micro-architecture in *Caenorhabditis elegans*": File S1

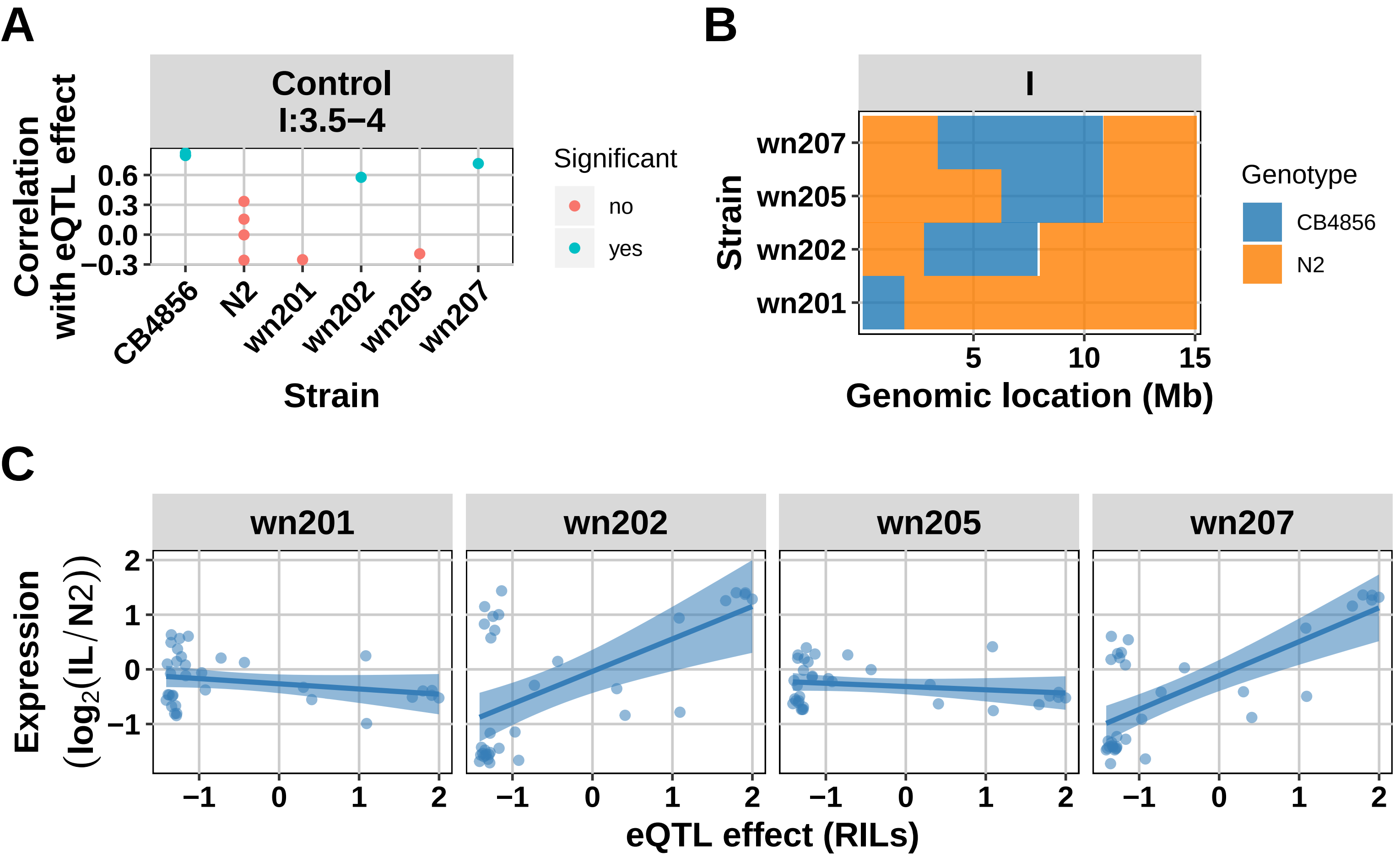
A
Control
I:3.5−4
Correlation with eQTL effect
0.6
0.3
0.0
−0.3
CB4856
N2
wn201
wn202
wn205
wn207
Strain
Significant
no
yes
B
I
wn207
wn205
wn202
wn201
Strain
5
10
15
Genomic location (Mb)
Genotype
CB4856
N2
C
wn201
wn202
wn205
wn207
Expression ($\log_2$(IL/N2))
2
1
0
−1
eQTL effect (RILs)

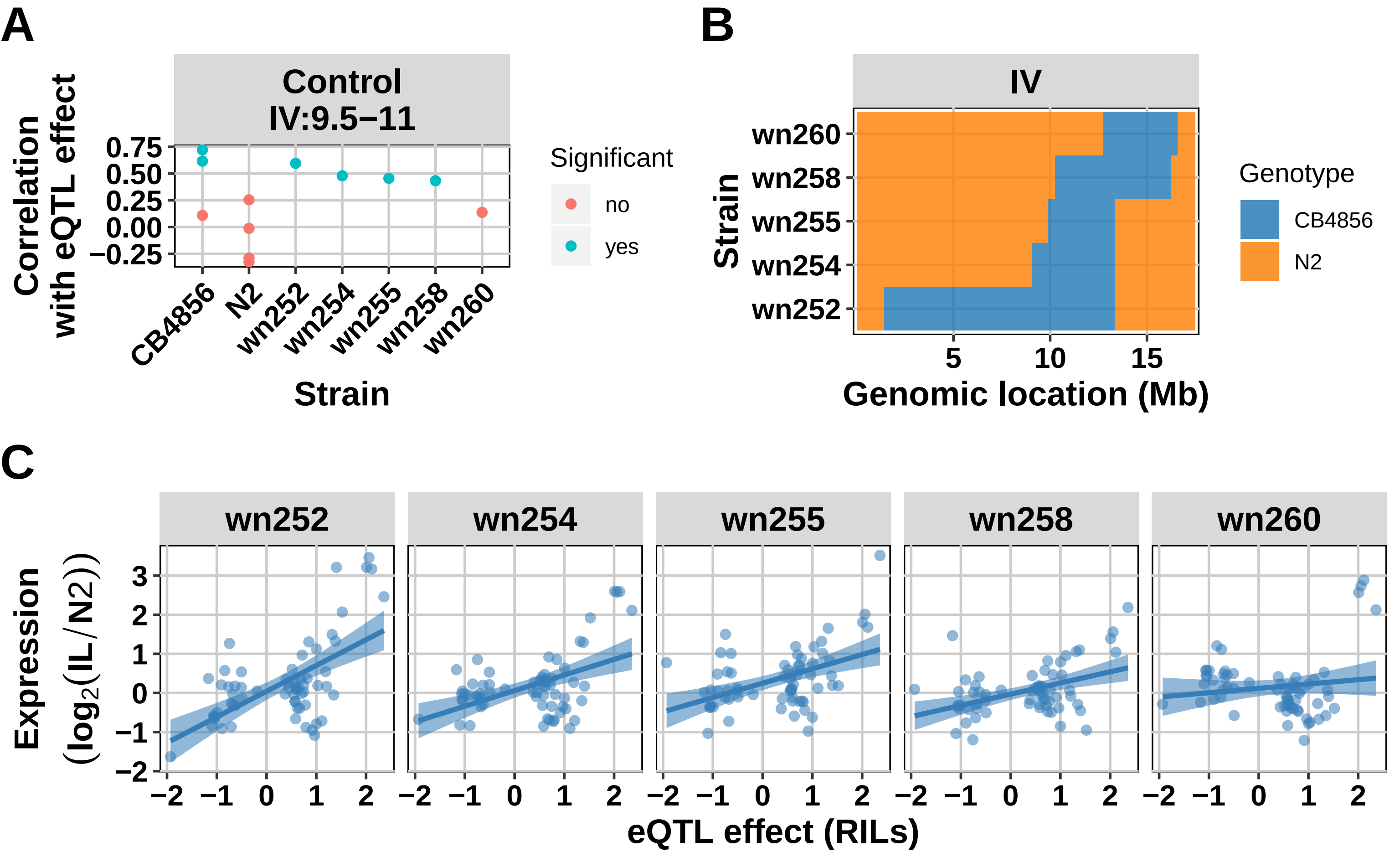

A
Control
IV:9.5−11
Correlation with eQTL effect
0.75
0.50
0.25
0.00
−0.25
CB4856
N2
wn252
wn254
wn255
wn258
wn260
Strain
Significant
no
yes
B
IV
wn260
wn258
wn255
wn254
wn252
Strain
5
10
15
Genomic location (Mb)
Genotype
CB4856
N2
C
wn252
wn254
wn255
wn258
wn260
Expression (log2(IL/N2))
3
2
1
0
−1
−2
−2 −1 0 1 2
eQTL effect (RILs)

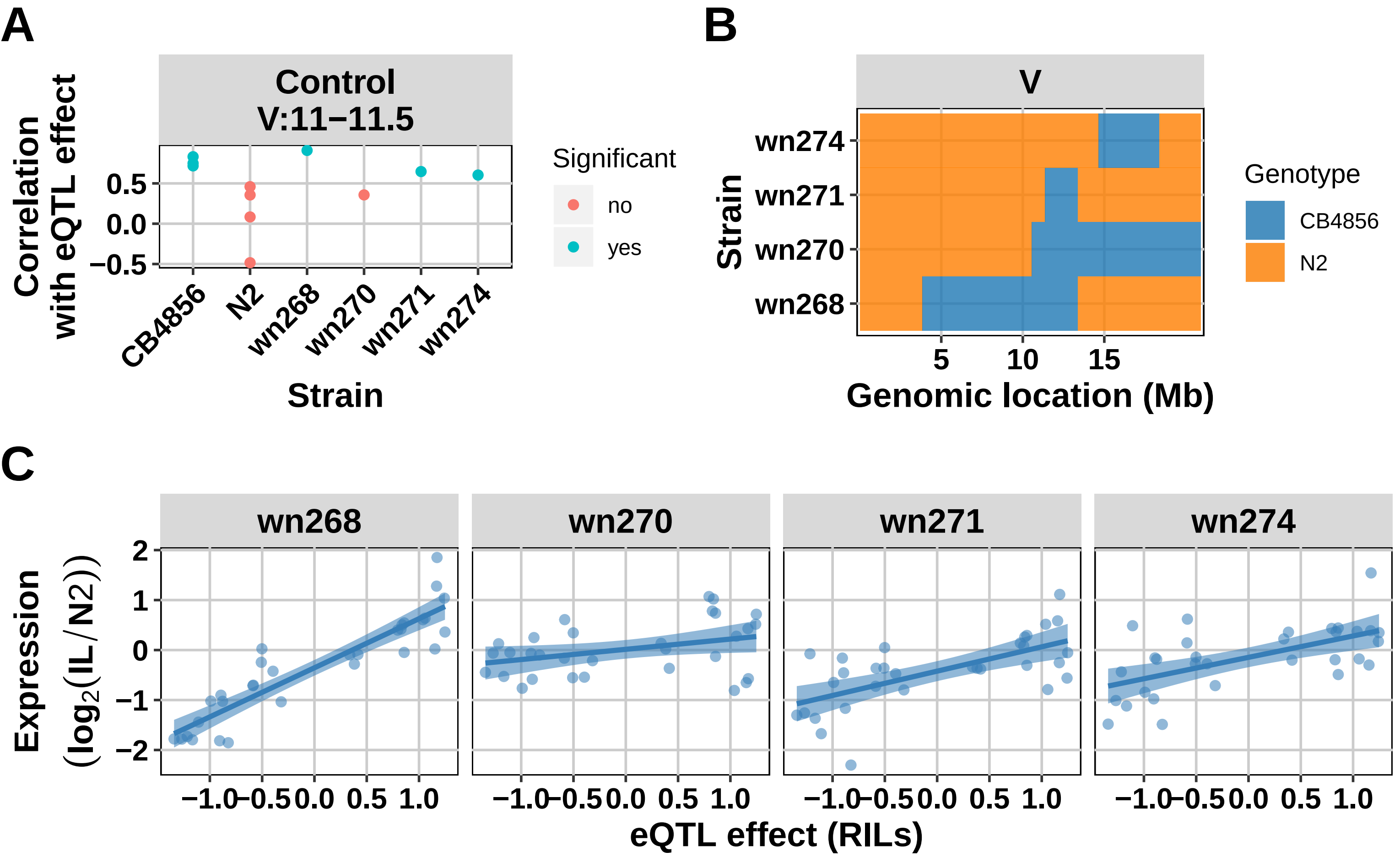

A
Control
V:11−11.5
Correlation with eQTL effect
0.5
0.0
−0.5
CB4856
N2
wn268
wn270
wn271
wn274
Strain
Significant
no
yes
B
V
wn274
wn271
wn270
wn268
Strain
5
10
15
Genomic location (Mb)
Genotype
CB4856
N2
C
wn268
wn270
wn271
wn274
Expression (log2(IL/N2))
2
1
0
−1
−2
−1.0
−0.5
0.0
0.5
1.0
eQTL effect (RILs)

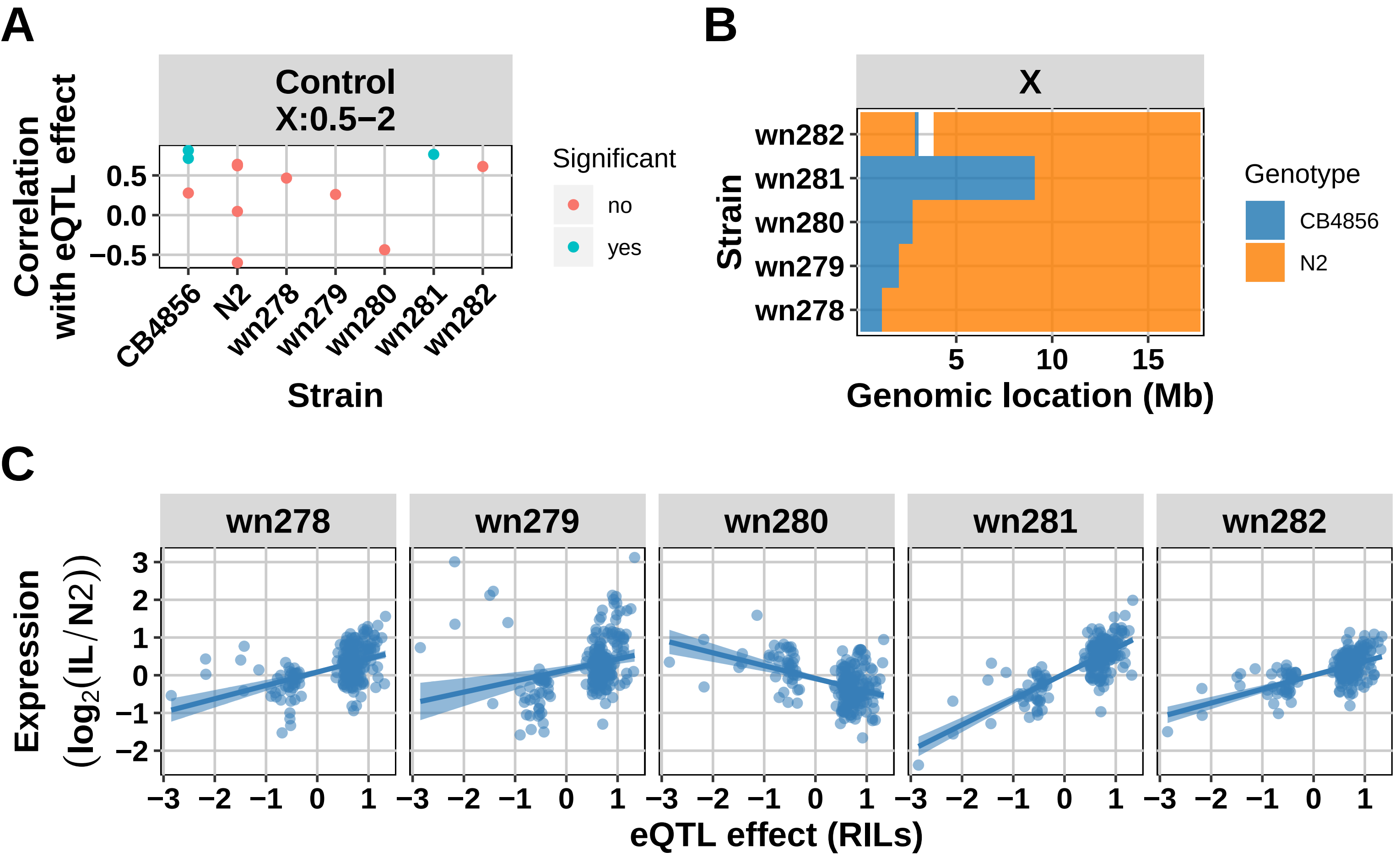

A
Control
X:0.5−2
Correlation with eQTL effect
0.5
0.0
−0.5
CB4856
N2
wn278
wn279
wn280
wn281
wn282
Strain
Significant
no
yes
B
X
wn282
wn281
wn280
wn279
wn278
Strain
5
10
15
Genomic location (Mb)
Genotype
CB4856
N2
C
wn278
wn279
wn280
wn281
wn282
Expression (log2(IL/N2))
3
2
1
0
−1
−2
−3
−2
−1
0
1
eQTL effect (RILs)

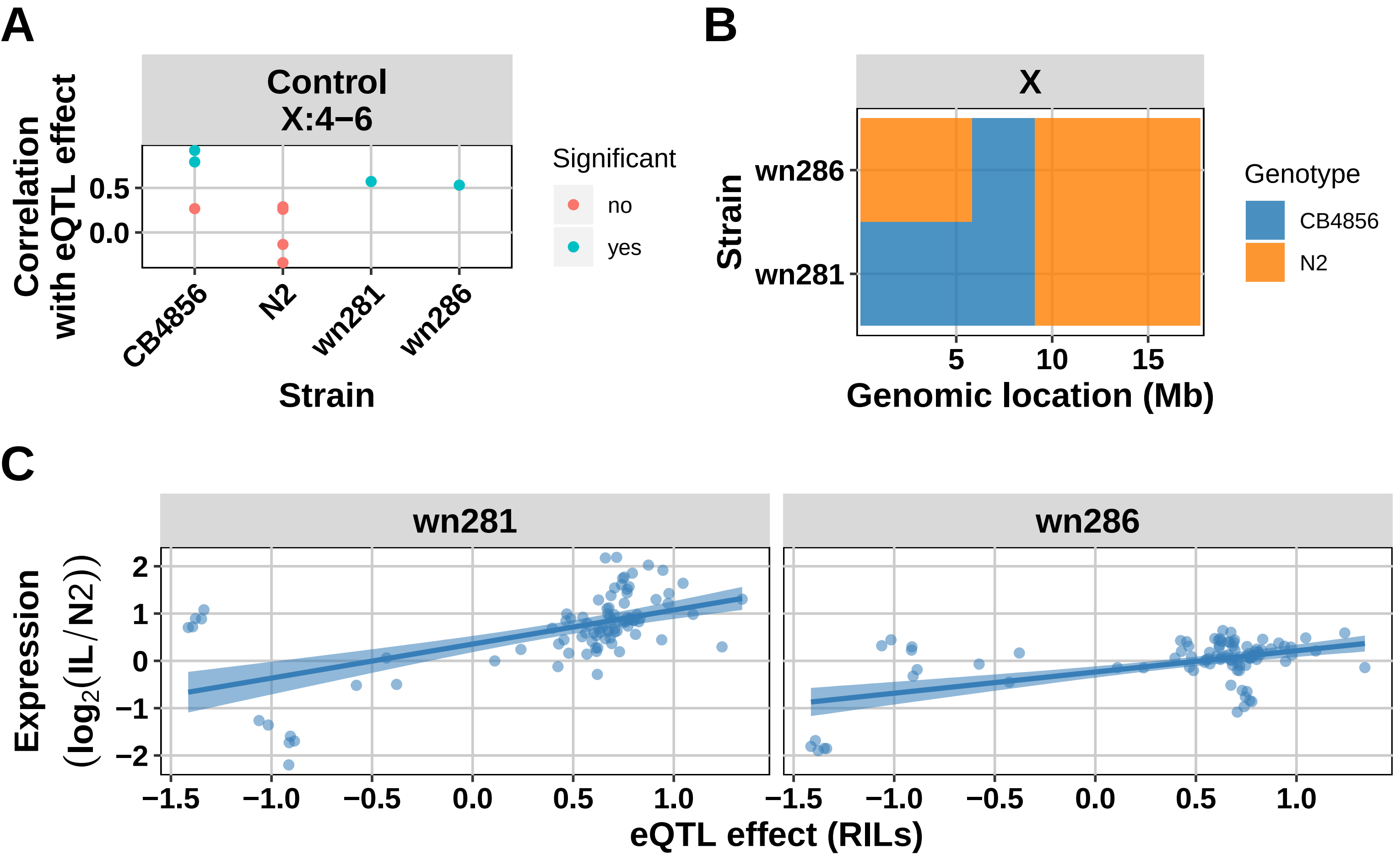
A
Control
X:4−6
Correlation with eQTL effect
0.5
0.0
CB4856
N2
wn281
wn286
Strain
Significant
no
yes
B
X
Strain
wn286
wn281
5
10
15
Genomic location (Mb)
Genotype
CB4856
N2
C
wn281
wn286
Expression ($\log_2$(IL/N2))
2
1
0
−1
−2
−1.5
−1.0
−0.5
0.0
0.5
1.0
eQTL effect (RILs)

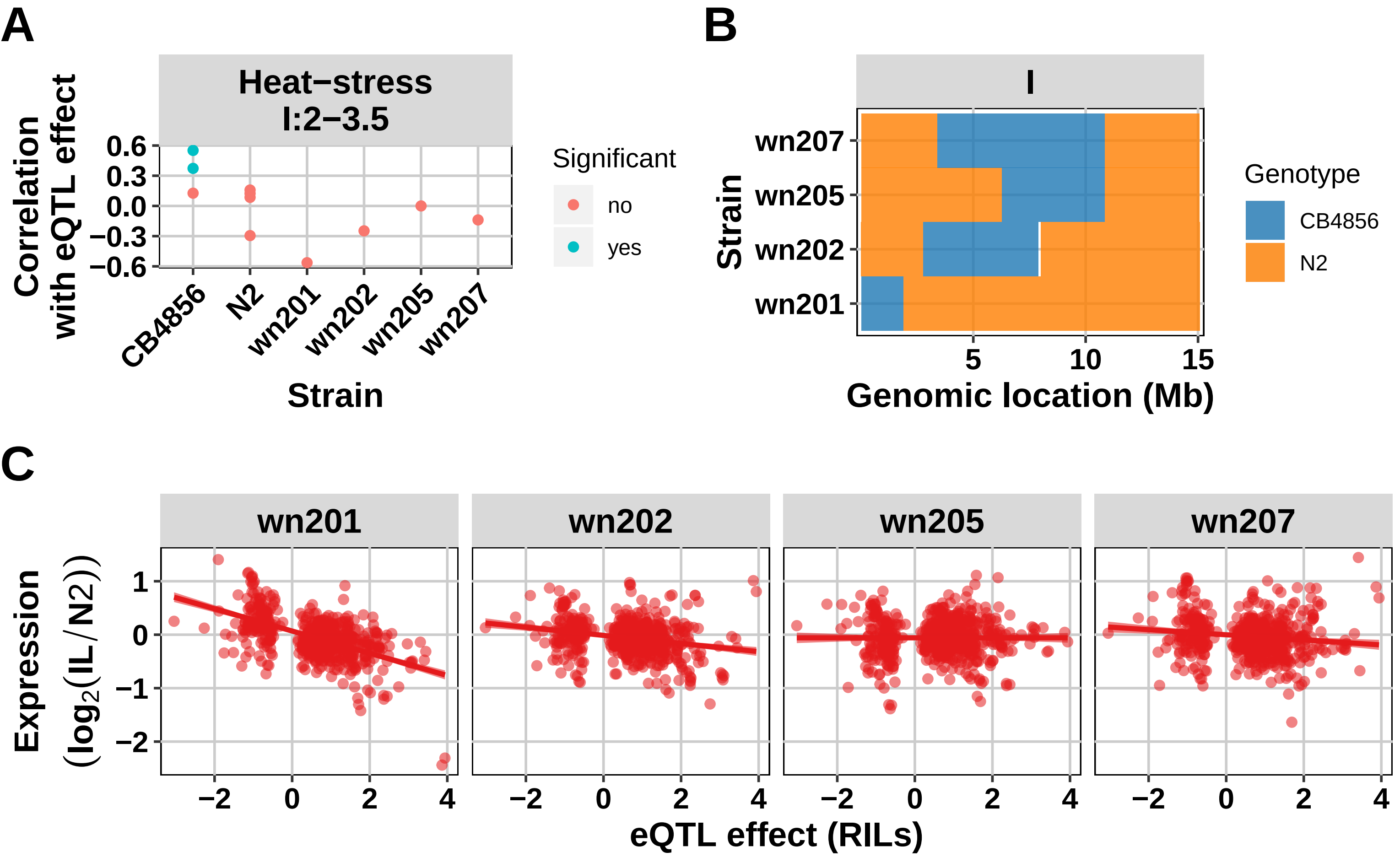
A
Heat−stress
I:2−3.5
Correlation with eQTL effect
0.6
0.3
0.0
−0.3
−0.6
CB4856
N2
wn201
wn202
wn205
wn207
Strain
Significant
no
yes
B
I
wn207
wn205
wn202
wn201
Strain
5
10
15
Genomic location (Mb)
Genotype
CB4856
N2
C
wn201
wn202
wn205
wn207
Expression (log2(IL/N2))
1
0
−1
−2
−2
0
2
4
eQTL effect (RILs)

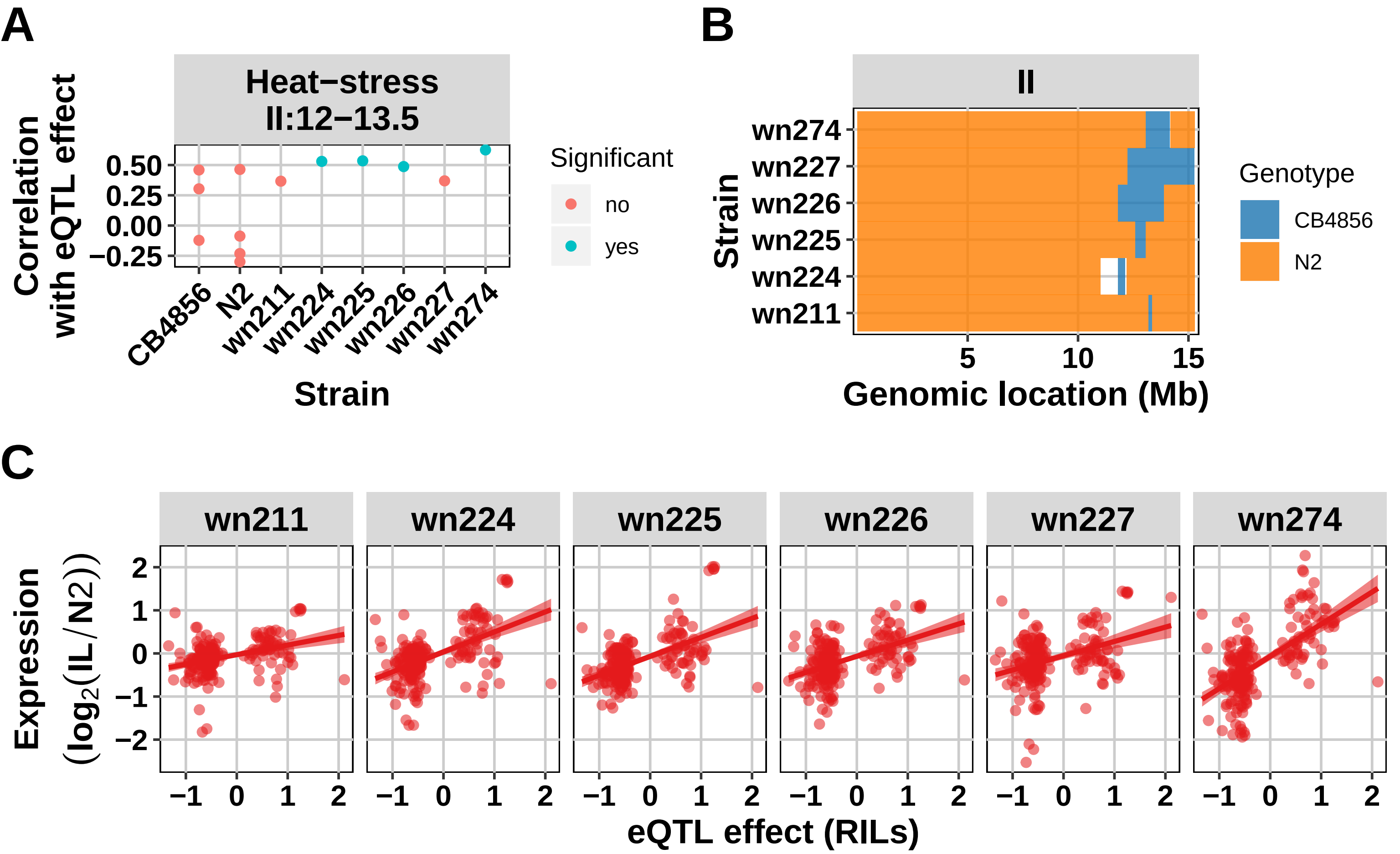
A
Heat−stress
II:12−13.5
Correlation with eQTL effect
0.50
0.25
0.00
−0.25
CB4856
N2
wn211
wn224
wn225
wn226
wn227
wn274
Strain
Significant
no
yes
B
II
wn274
wn227
wn226
wn225
wn224
wn211
Strain
5
10
15
Genomic location (Mb)
Genotype
CB4856
N2
C
wn211
wn224
wn225
wn226
wn227
wn274
Expression (log2(IL/N2))
2
1
0
−1
−2
−1
0
1
2
eQTL effect (RILs)

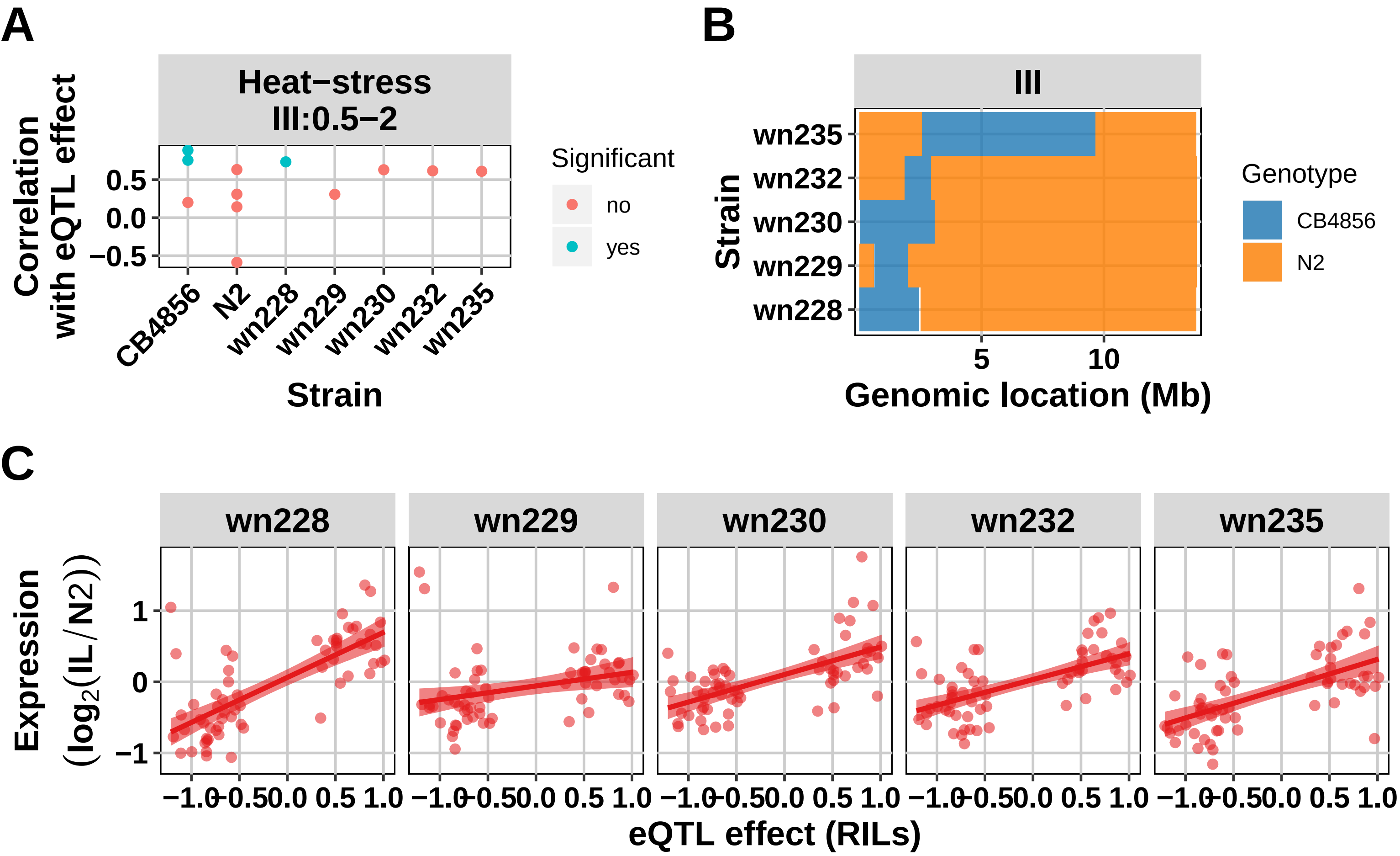
A
Heat−stress
III:0.5−2
Correlation with eQTL effect
0.5
0.0
−0.5
CB4856
N2
wn228
wn229
wn230
wn232
wn235
Strain
Significant
no
yes
B
III
Strain
wn235
wn232
wn230
wn229
wn228
5
10
Genomic location (Mb)
Genotype
CB4856
N2
C
wn228
wn229
wn230
wn232
wn235
Expression (log2(IL/N2))
1
0
−1
−1.0
−0.5
0.0
0.5
1.0
eQTL effect (RILs)

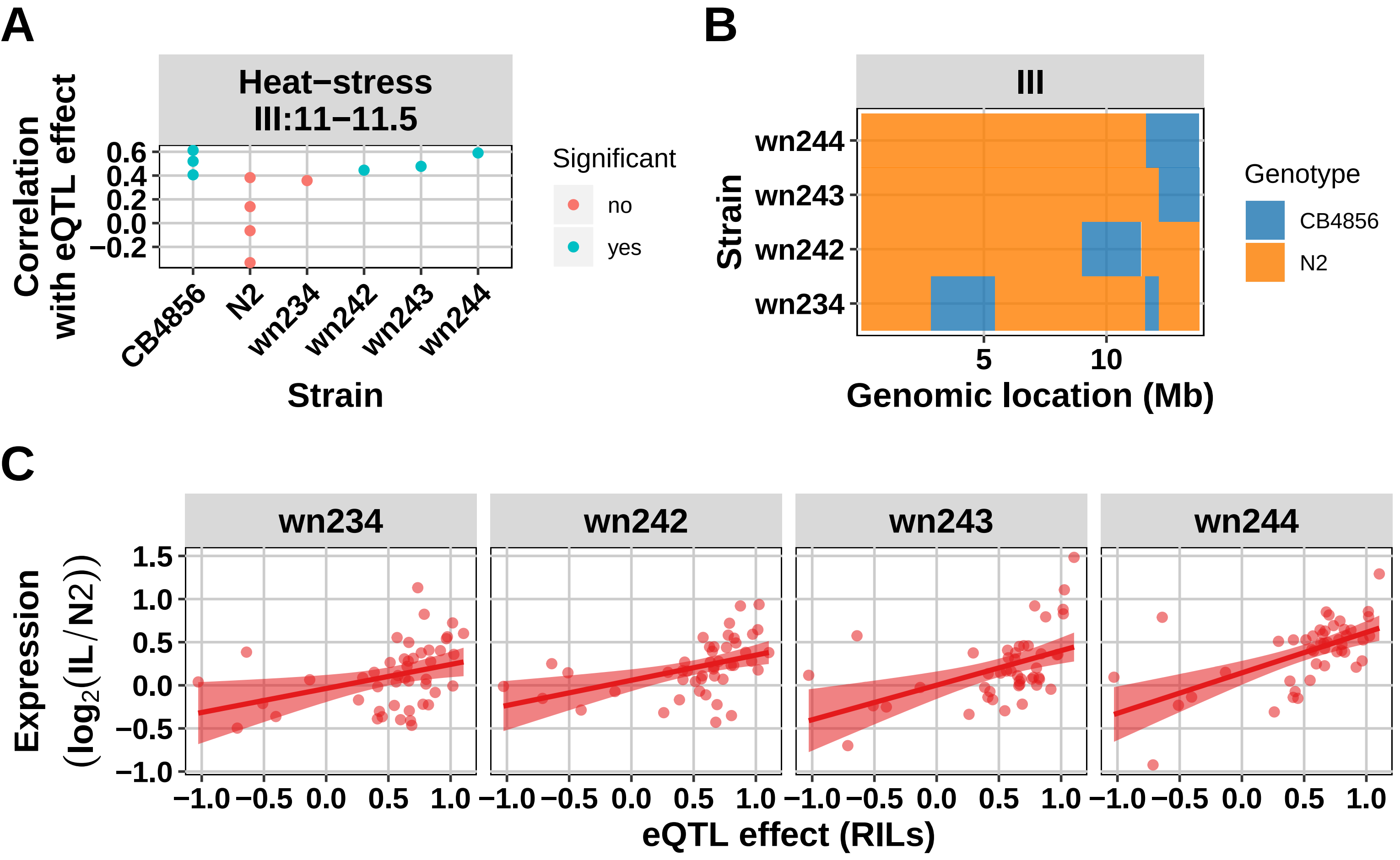
A
Heat−stress
III:11−11.5
Correlation with eQTL effect
0.6
0.4
0.2
0.0
−0.2
CB4856
N2
wn234
wn242
wn243
wn244
Strain
Significant
no
yes
B
III
wn244
wn243
wn242
wn234
Strain
5
10
Genomic location (Mb)
Genotype
CB4856
N2
C
wn234
wn242
wn243
wn244
1.5
1.0
0.5
0.0
−0.5
−1.0
Expression (log2(IL/N2))
−1.0
−0.5
0.0
0.5
1.0
eQTL effect (RILs)

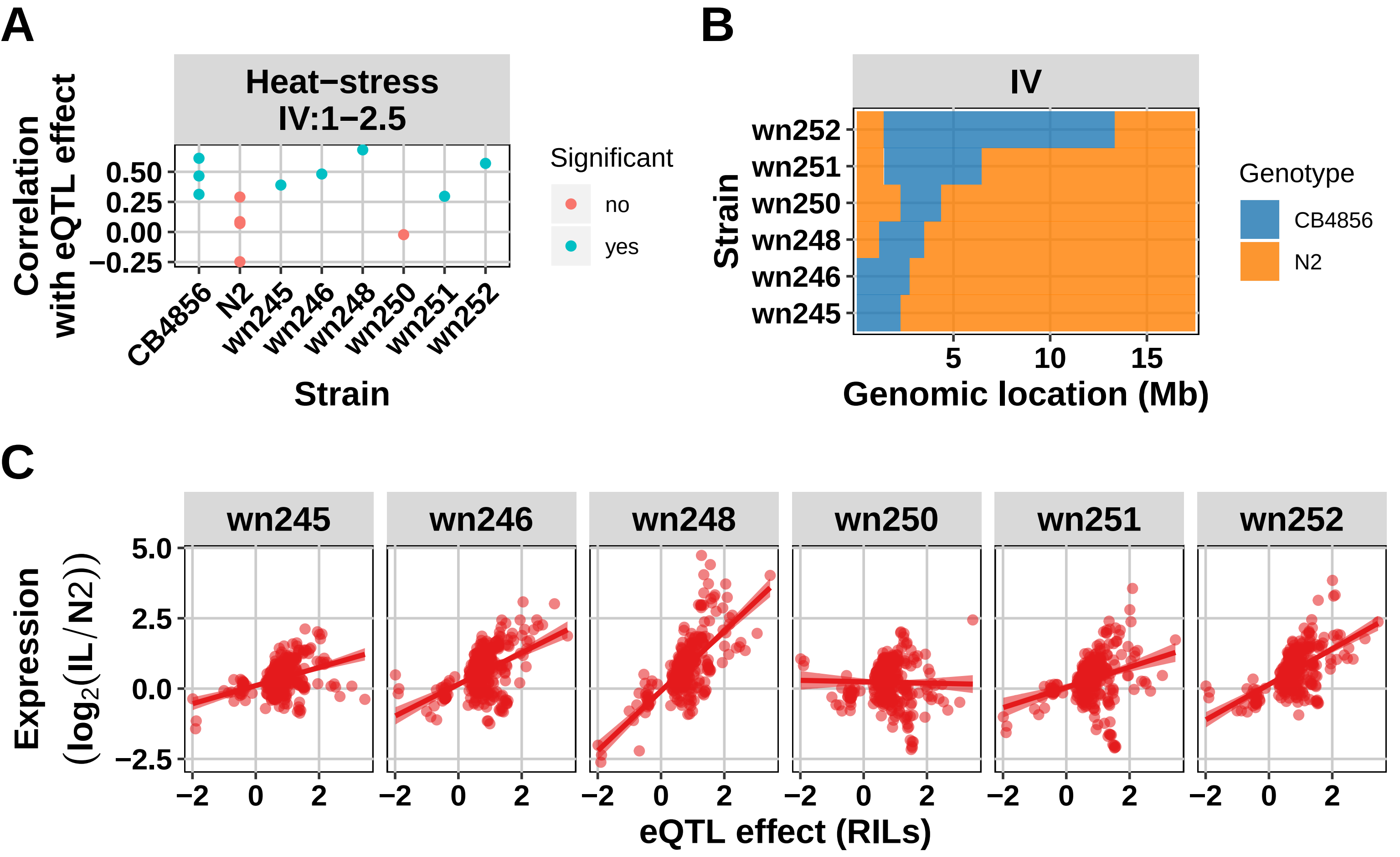

A
Heat−stress
IV:1−2.5
Correlation with eQTL effect
0.50
0.25
0.00
−0.25
CB4856
N2
wn245
wn246
wn248
wn250
wn251
wn252
Strain
Significant
no
yes
B
IV
Strain
wn252
wn251
wn250
wn248
wn246
wn245
5
10
15
Genomic location (Mb)
Genotype
CB4856
N2
C
wn245
wn246
wn248
wn250
wn251
wn252
Expression (log2(IL/N2))
5.0
2.5
0.0
−2.5
−2
0
2
eQTL effect (RILs)

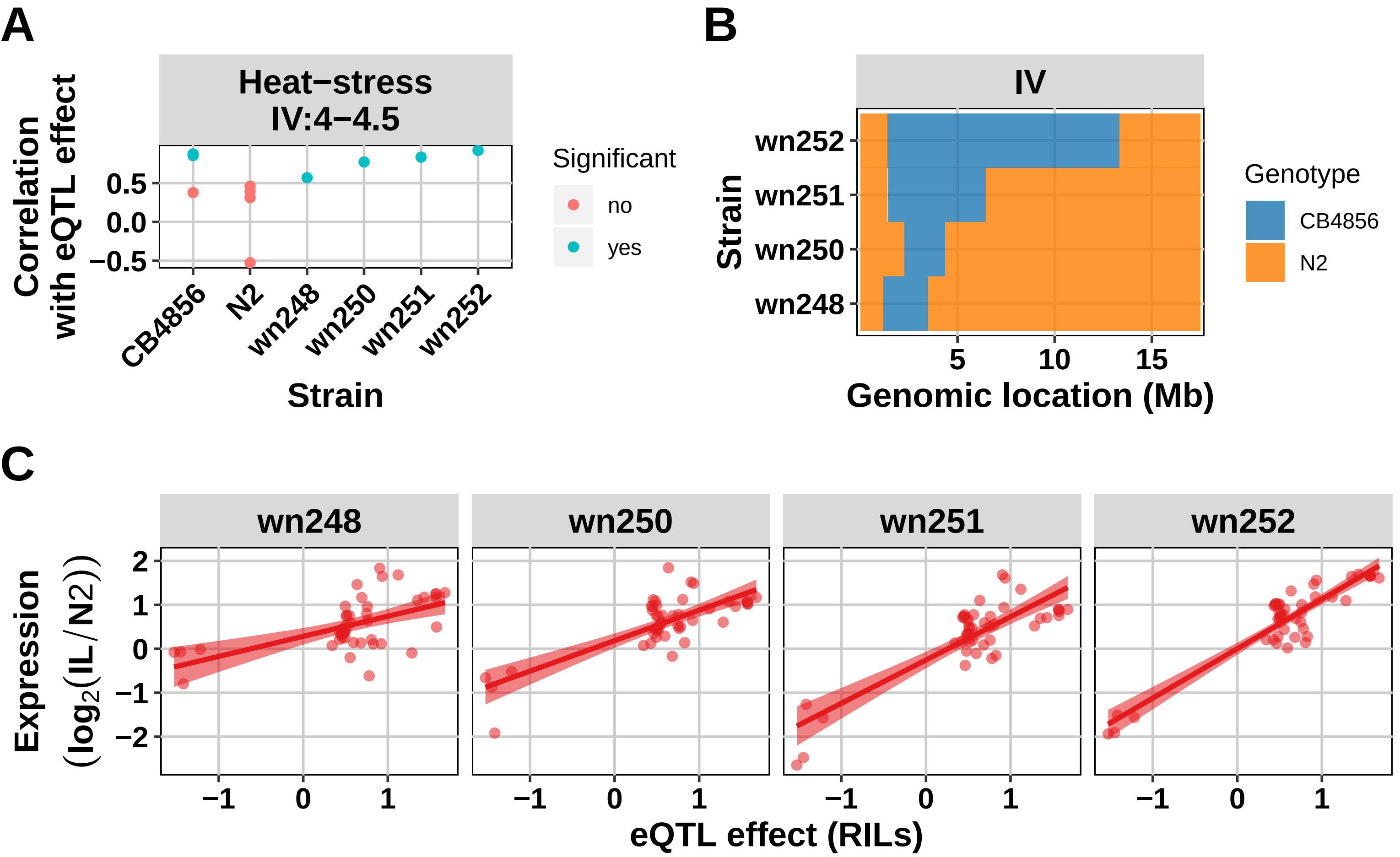
A
Heat−stress
IV:4−4.5
Correlation with eQTL effect
0.5
0.0
−0.5
CB4856
N2
wn248
wn250
wn251
wn252
Strain
Significant
no
yes
B
IV
wn252
wn251
wn250
wn248
Strain
5
10
15
Genomic location (Mb)
Genotype
CB4856
N2
C
wn248
wn250
wn251
wn252
Expression ($log_2$(IL/N2))
2
1
0
−1
−2
−1
0
1
eQTL effect (RILs)

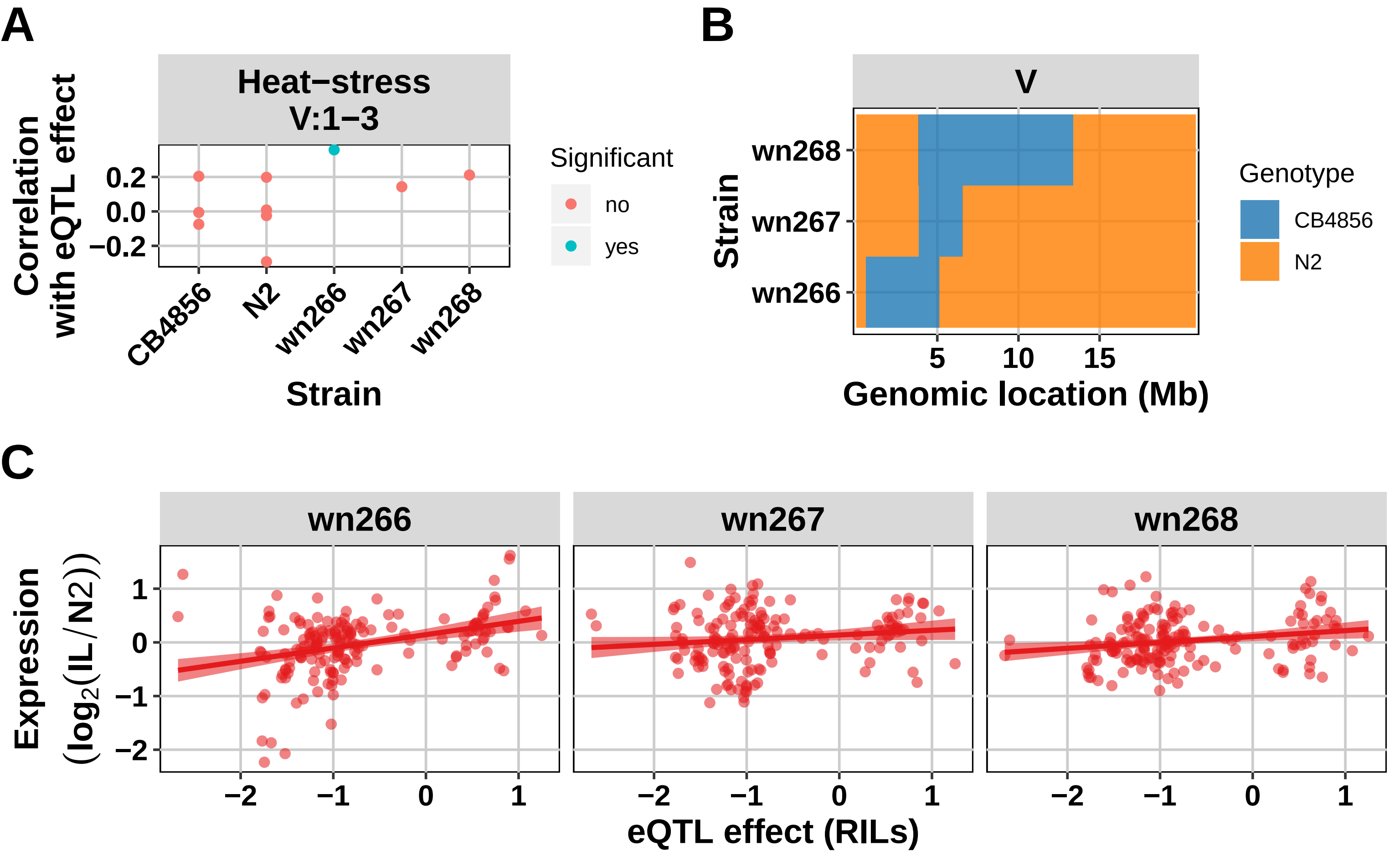

A
Heat−stress
V:1−3
Correlation with eQTL effect
0.2
0.0
−0.2
CB4856
N2
wn266
wn267
wn268
Strain
Significant
no
yes
B
V
wn268
wn267
wn266
Strain
5
10
15
Genomic location (Mb)
Genotype
CB4856
N2
C
wn266
wn267
wn268
Expression (log2(IL/N2))
1
0
−1
−2
−2
−1
0
1
eQTL effect (RILs)

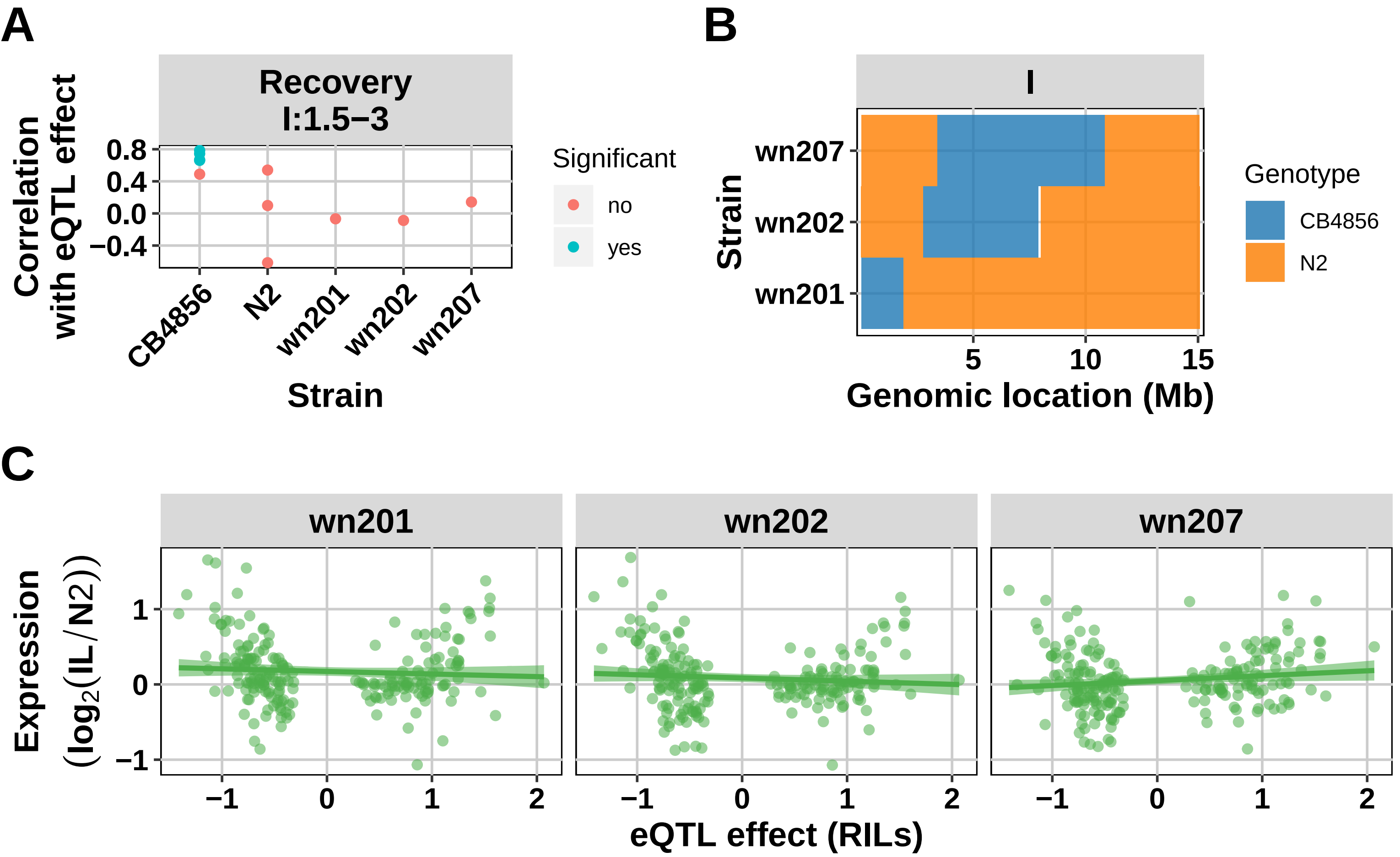
A
Recovery
I:1.5–3
Correlation with eQTL effect
0.8
0.4
0.0
−0.4
CB4856
N2
wn201
wn202
wn207
Strain
Significant
no
yes
B
I
Strain
wn207
wn202
wn201
5
10
15
Genomic location (Mb)
Genotype
CB4856
N2
C
wn201
wn202
wn207
Expression (log2(IL/N2))
1
0
−1
−1
0
1
2
eQTL effect (RILs)

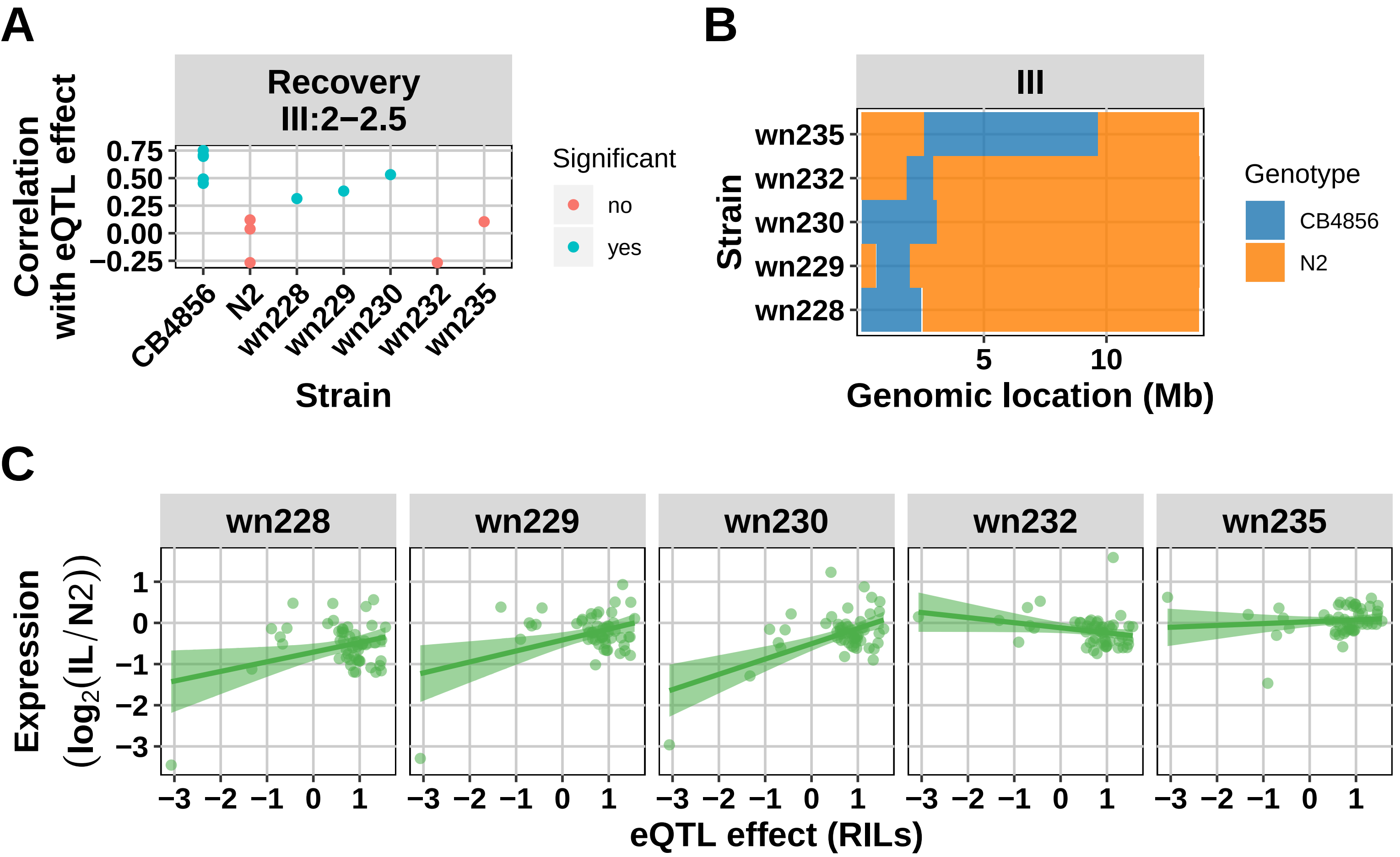

A
Recovery
III:2−2.5
Correlation with eQTL effect
0.75
0.50
0.25
0.00
−0.25
CB4856
N2
wn228
wn229
wn230
wn232
wn235
Strain
Significant
no
yes
B
III
wn235
wn232
wn230
wn229
wn228
Strain
5
10
Genomic location (Mb)
Genotype
CB4856
N2
C
wn228
wn229
wn230
wn232
wn235
Expression (log2(IL/N2))
1
0
−1
−2
−3
eQTL effect (RILs)

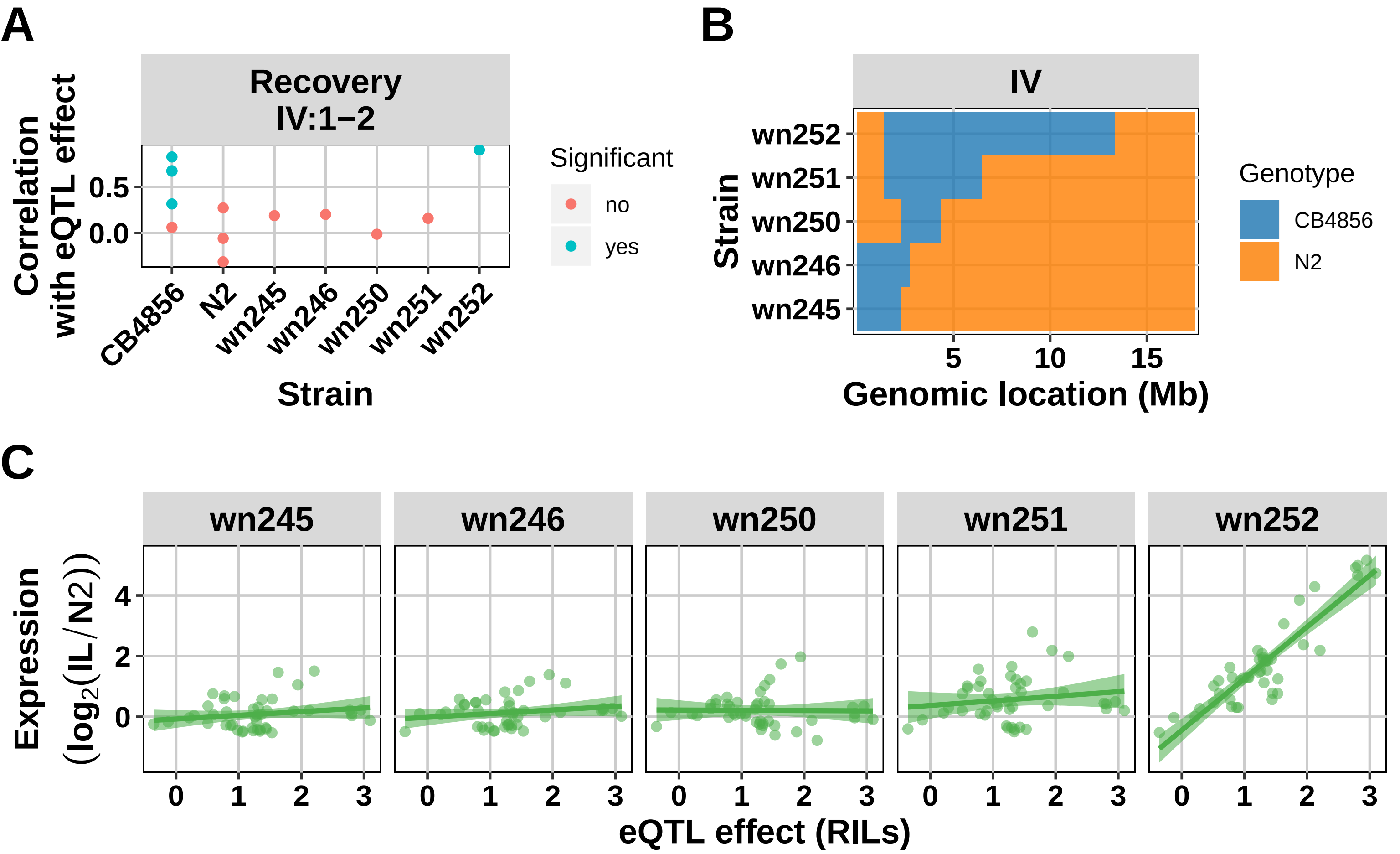
A
Recovery
IV:1−2
Correlation with eQTL effect
0.5
0.0
CB4856
N2
wn245
wn246
wn250
wn251
wn252
Strain
Significant
no
yes
B
IV
wn252
wn251
wn250
wn246
wn245
Strain
5
10
15
Genomic location (Mb)
Genotype
CB4856
N2
C
wn245
wn246
wn250
wn251
wn252
Expression (log2(IL/N2))
4
2
0
0
1
2
3
eQTL effect (RILs)

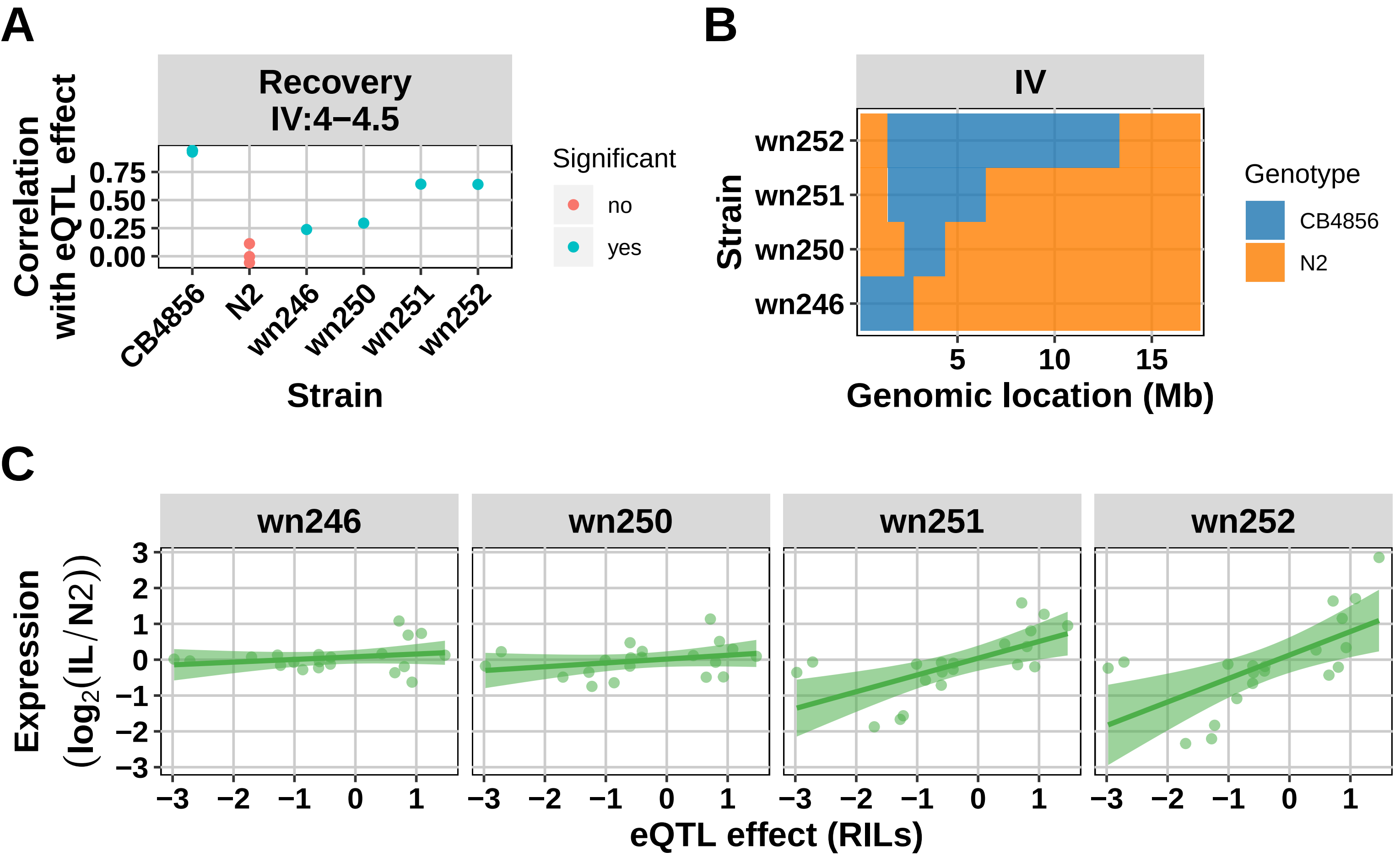

A
Recovery
IV:4−4.5
Correlation with eQTL effect
0.75
0.50
0.25
0.00
CB4856
N2
wn246
wn250
wn251
wn252
Strain
Significant
no
yes
B
IV
wn252
wn251
wn250
wn246
Strain
5
10
15
Genomic location (Mb)
Genotype
CB4856
N2
C
wn246
wn250
wn251
wn252
Expression (log2(IL/N2))
3
2
1
0
−1
−2
−3
−3
−2
−1
0
1
eQTL effect (RILs)

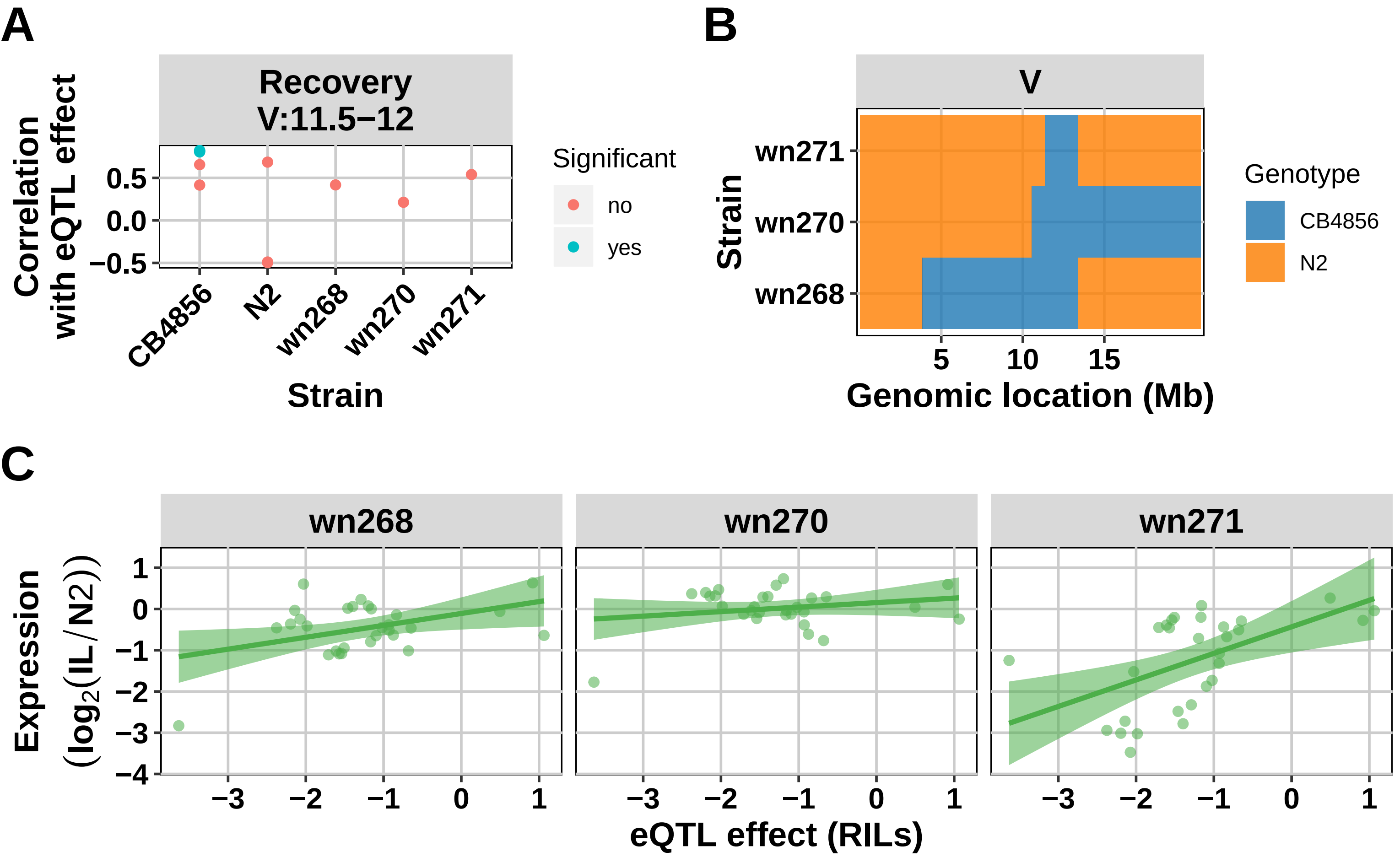

A
Recovery
V:11.5−12
Correlation with eQTL effect
0.5
0.0
−0.5
CB4856
N2
wn268
wn270
wn271
Strain
Significant
no
yes
B
V
wn271
wn270
wn268
Strain
5
10
15
Genomic location (Mb)
Genotype
CB4856
N2
C
wn268
wn270
wn271
Expression (log2(IL/N2))
1
0
−1
−2
−3
−4
−3
−2
−1
0
1
eQTL effect (RILs)

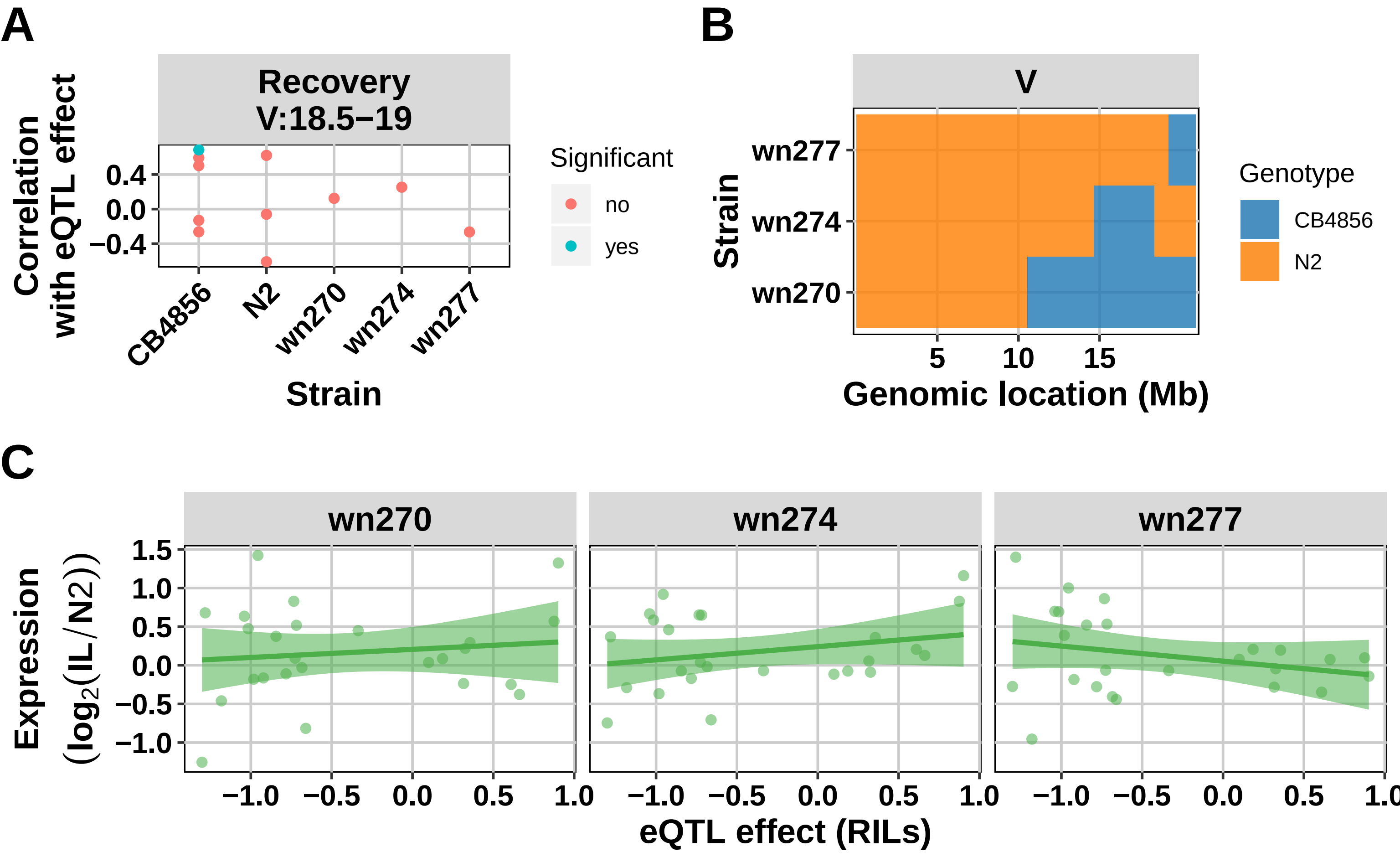
A
Recovery
V:18.5−19
Correlation with eQTL effect
0.4
0.0
−0.4
CB4856
N2
wn270
wn274
wn277
Strain
Significant
no
yes
B
V
Strain
wn277
wn274
wn270
5
10
15
Genomic location (Mb)
Genotype
CB4856
N2
C
wn270
wn274
wn277
Expression (log2(IL/N2))
1.5
1.0
0.5
0.0
−0.5
−1.0
−1.0
−0.5
0.0
0.5
1.0
eQTL effect (RILs)

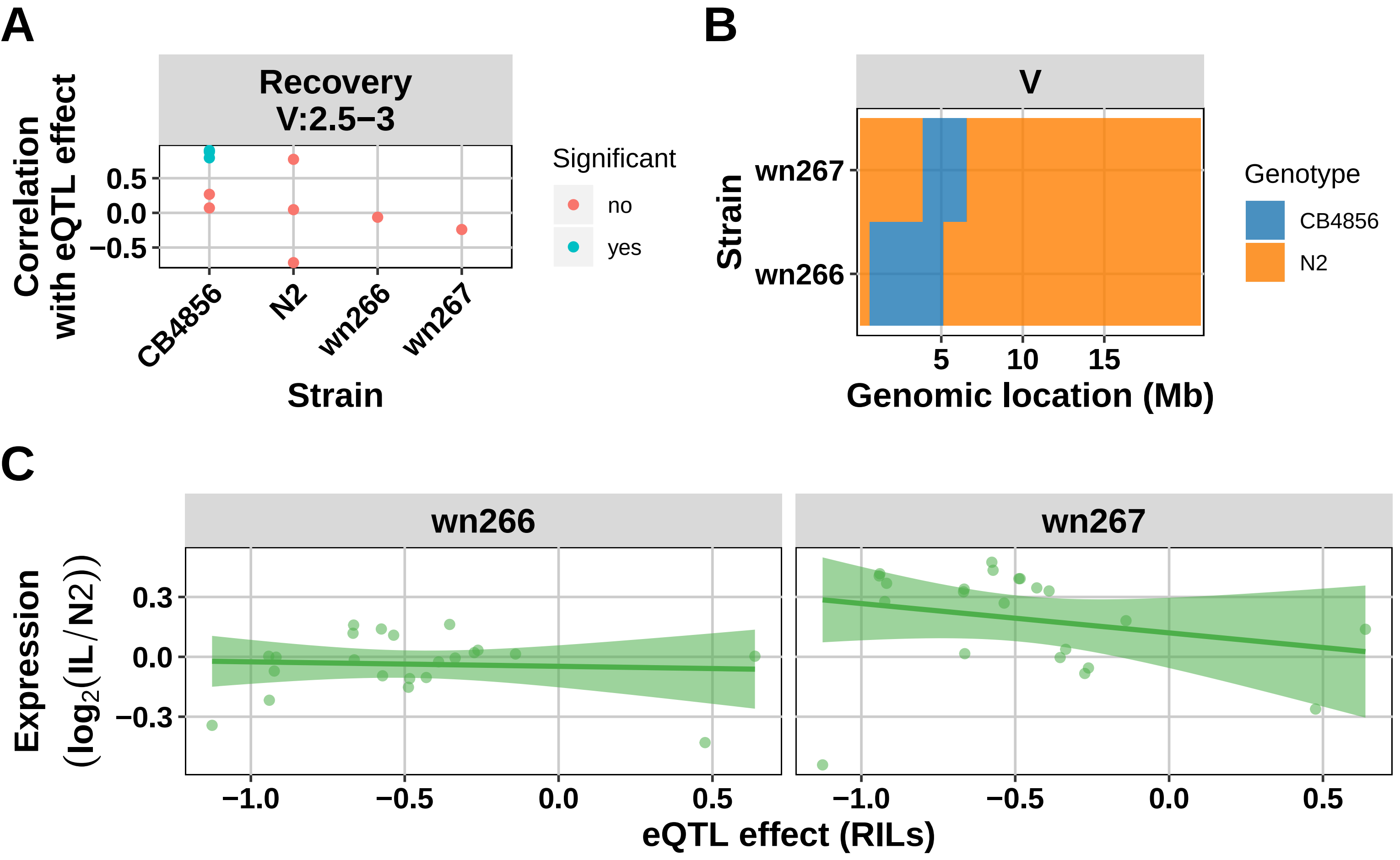

A
Recovery
V:2.5−3
Correlation with eQTL effect
0.5
0.0
−0.5
CB4856
N2
wn266
wn267
Strain
Significant
no
yes
B
V
wn267
wn266
Strain
5
10
15
Genomic location (Mb)
Genotype
CB4856
N2
C
wn266
wn267
Expression (log2(IL/N2))
0.3
0.0
−0.3
−1.0
−0.5
0.0
0.5
eQTL effect (RILs)
