## Supplementary figures and images for "Dissecting the eQTL micro-architecture in *Caenorhabditis elegans*"

### Figure S1

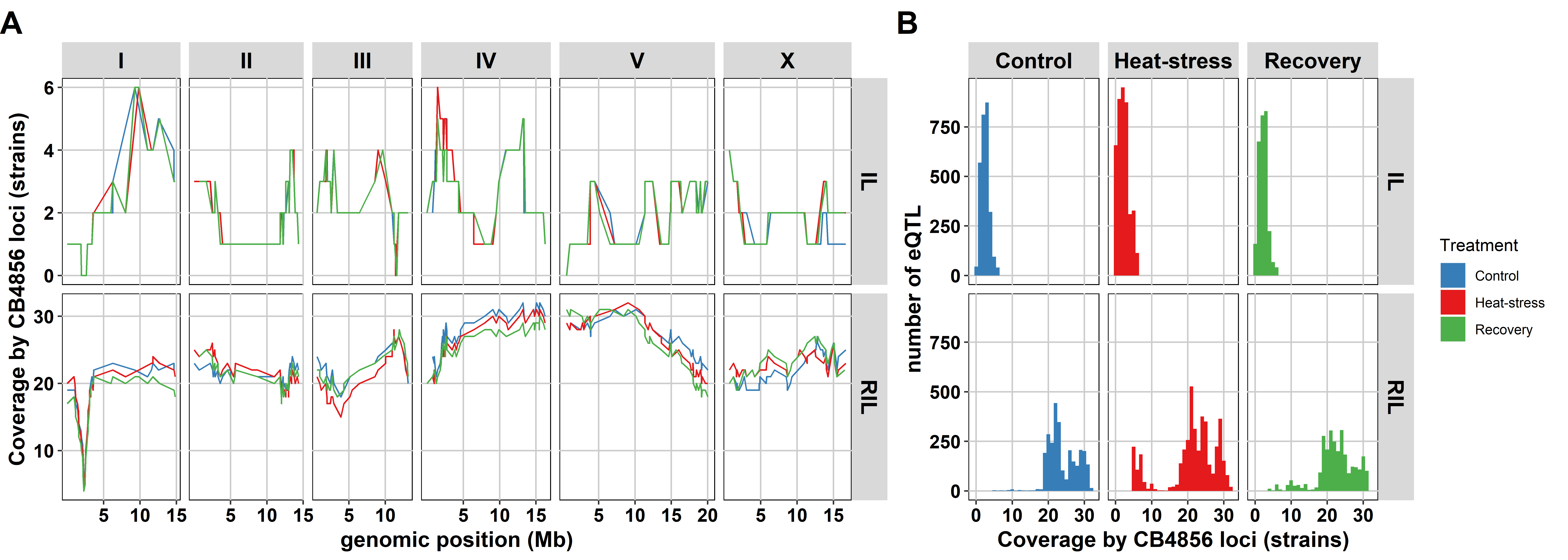

### Figure S2

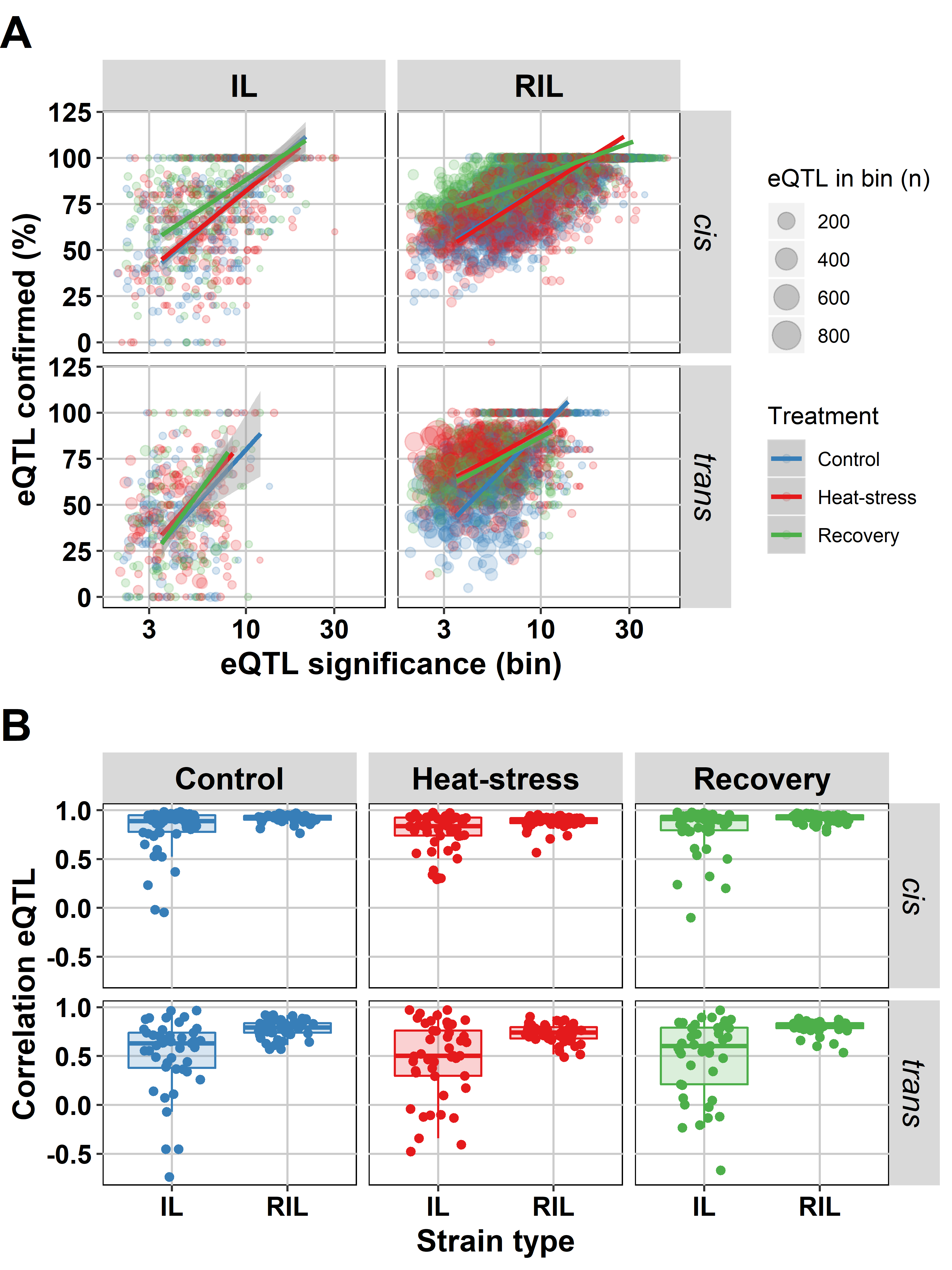

### Figure S3

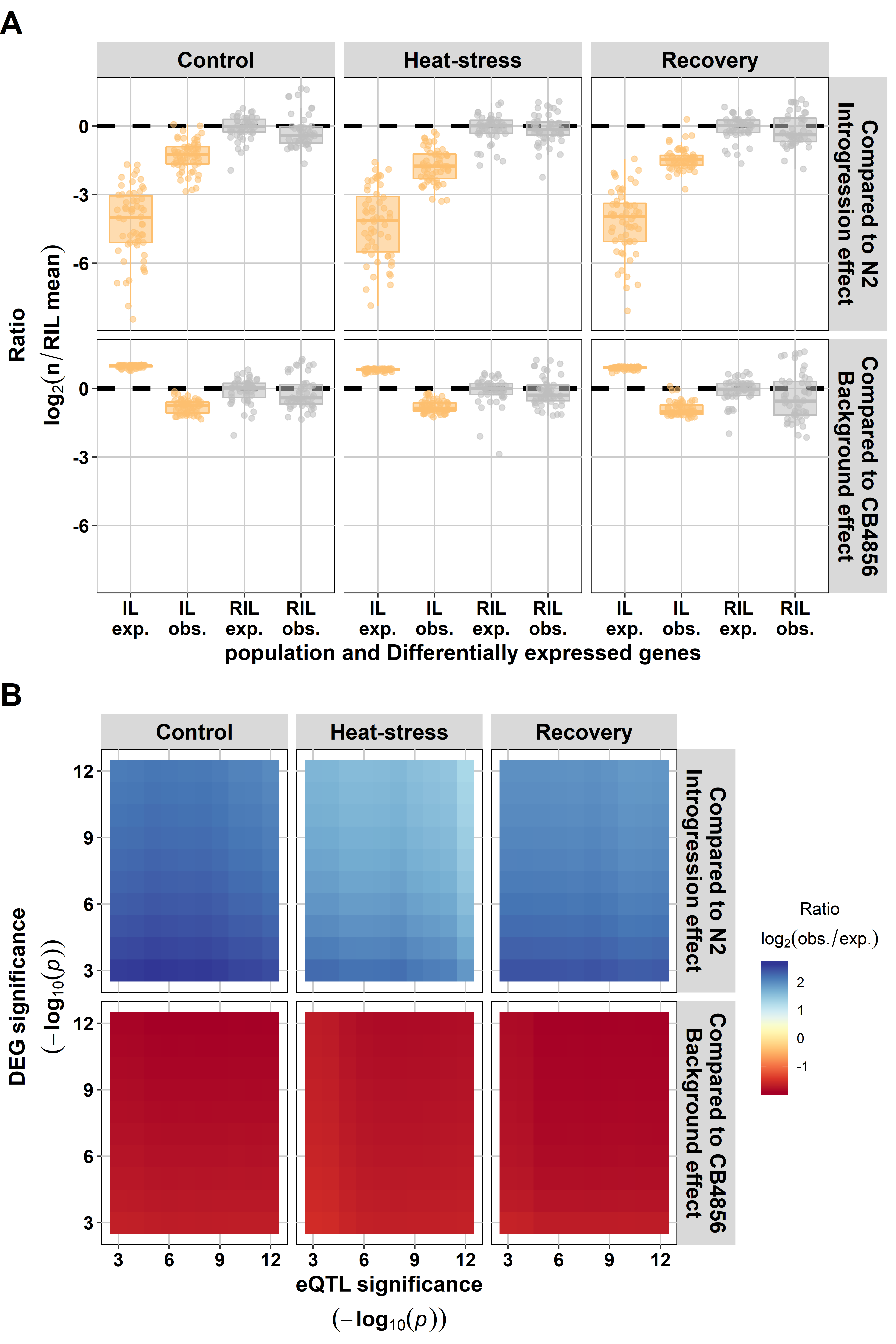

### Figure S4

#### Slide 1
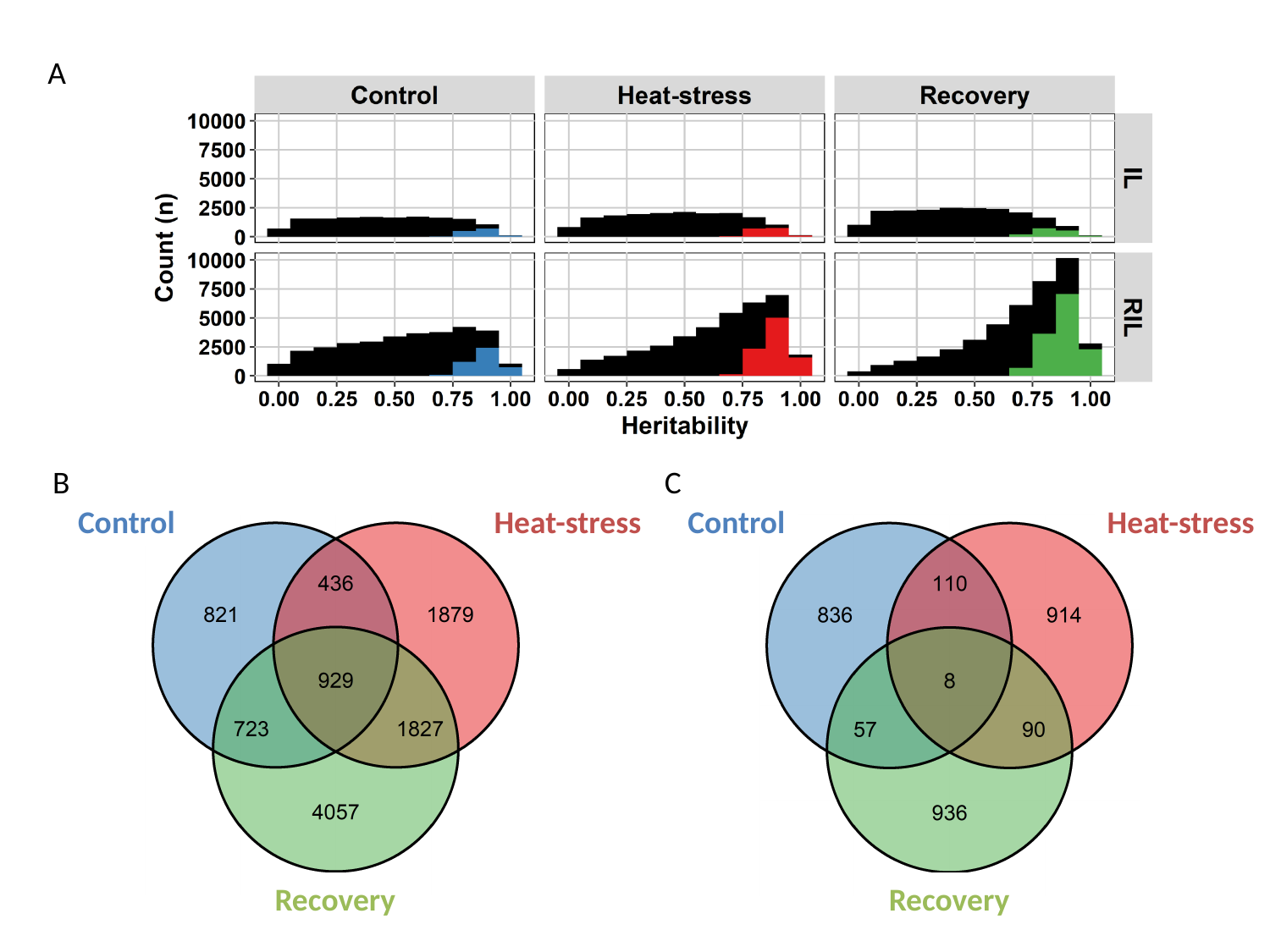

A
B
C
Control
Heat-stress
Control
Heat-stress
Recovery
Recovery

### Figure S6

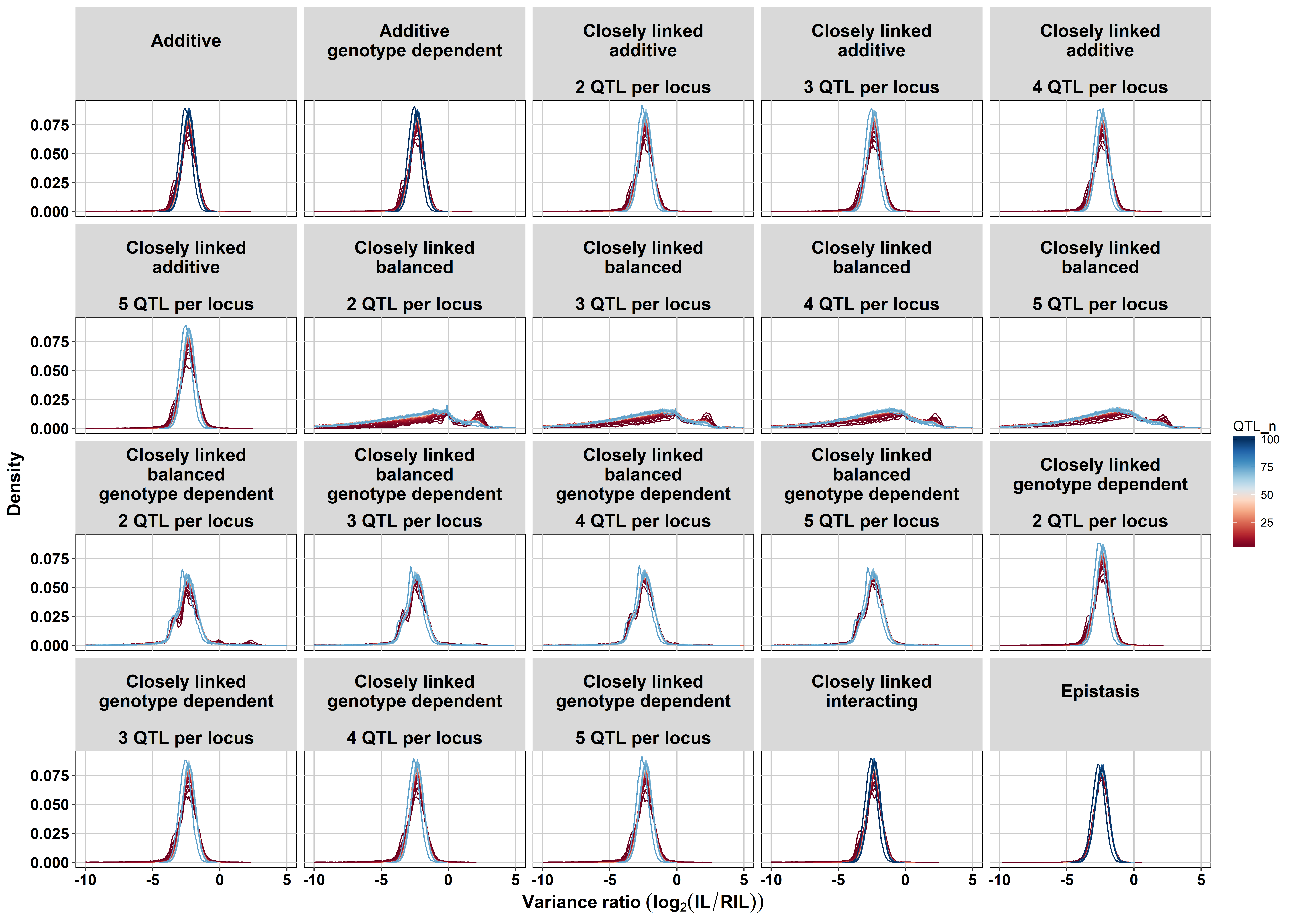

### Figure S7

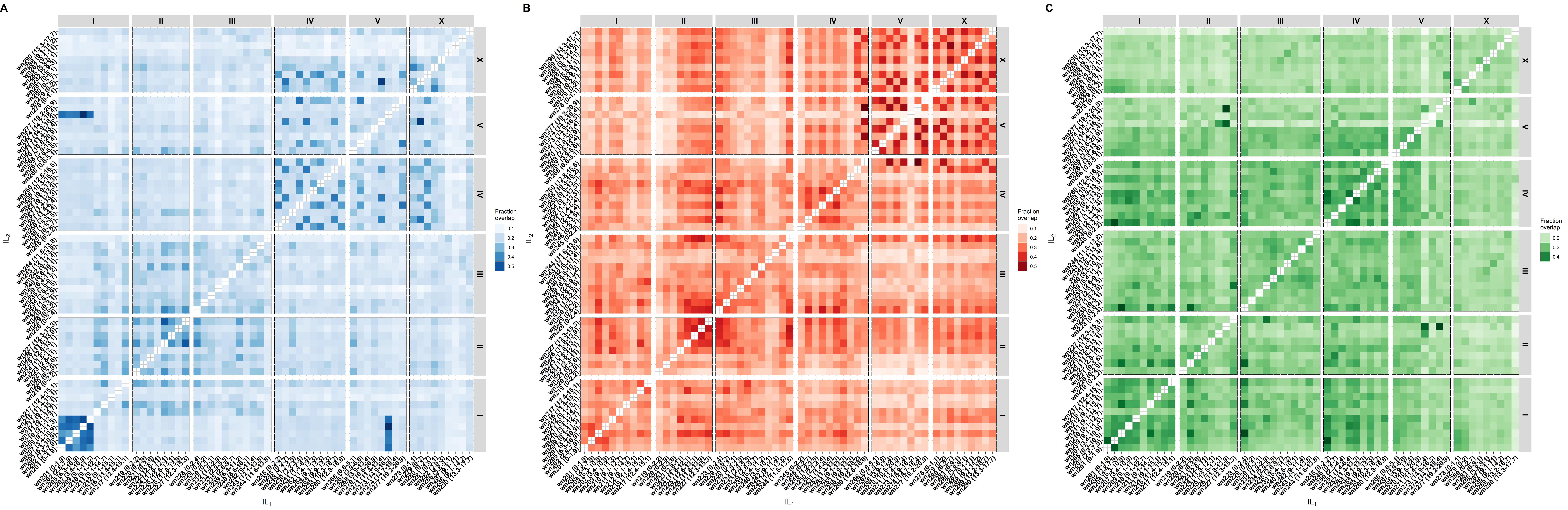
