## Supplementary material for "Dissecting the eQTL micro-architecture in *Caenorhabditis elegans*": Figure S5

### Slide 1
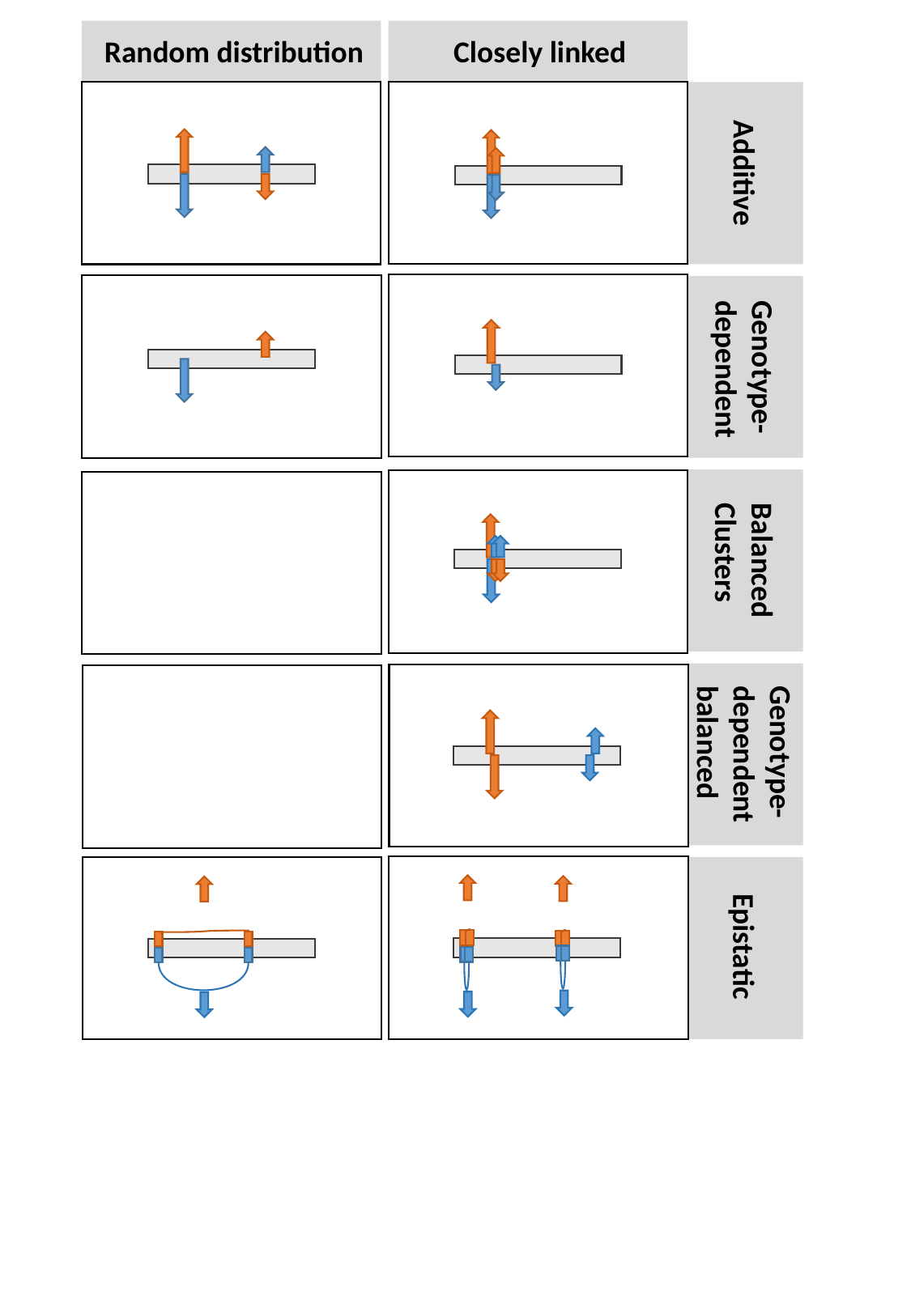

Closely linked
Random distribution
Additive
Genotype-
dependent
Balanced
Clusters
Genotype-
dependent
balanced
Epistatic
